# RNA Structure Coordinates Translation Across the Meiotic Program

**DOI:** 10.64898/2026.01.15.699792

**Authors:** Hao Wu, Caini Zhou, Shoucheng Du, Simon Felder, Yihan Xiong, Kevin M. Weeks, Yun Bai

**Affiliations:** School of Life Science and Technology, ShanghaiTech University, 201210 Shanghai, China; Department of Chemistry, University of North Carolina, Chapel Hill, NC 27599-3290, USA

**Keywords:** RNA structure, meiosis, translation timing, RNA helicase

## Abstract

mRNA structure clearly modulates translation efficiency for individual transcripts, yet its role in coordinating translation across many transcripts remains poorly defined. Meiosis offers a compelling test case, involving large-scale gene expression changes coordinated across hundreds of mRNAs under conditions where transcription is constrained by chromosome condensation. We profiled mRNA structures across yeast meiosis, generating a high-resolution structurome encompassing ∼70% of annotated mRNAs, including multi-time-point measurements for 2,084 transcripts. Transcripts upregulated during meiosis generally adopt flexible structures that enhance translation, whereas mRNAs with complex 5′ UTR and coding-region structures show suppressed translation, indicating that RNA structure globally shapes meiotic translation. We further observed a high-low-high oscillation in cytoplasmic RNA helicase levels across meiosis. Together, these findings support a model in which RNA structure and helicase expression act jointly to program stage-specific translation of hundreds of mRNAs, with highly structured transcripts selectively translated at late stages. Consistent with this model, disrupting the structure of the abundant transcript *CCW22*, or altering Ded1p helicase levels, impaired meiotic progression in an RNA structure-dependent manner. Our study reveals that the concerted action of RNA structure and helicases coordinates cell-wide translation dynamics to meet stage-specific demands, especially critical when transcription is limited during meiotic divisions.

## INTRODUCTION

Elaborate structures in RNA molecules have diverse regulatory functions^1^. High-throughput structure-probing has revealed RNA structural landscapes in diverse entities, from viruses to animals^2–7^. Foundational work has shown that RNA structure can have switch-like functions that modulate the expression of individual mRNAs in dynamic cellular contexts, including isolated cellular compartments^8^, different phases of zebrafish embryogenesis^9^, and pattern-triggered immunity in *Arabidopsis*^10^. However, the substantial sequencing depth required for reliable structure probing has historically limited transcriptome-wide coverage, making it difficult to uncover general rules for how RNA structure globally coordinates gene expression during dynamic cellular processes.

RNA structure profoundly impacts translation, a process inherently dependent on mRNA-ribosome interactions and structural barriers. Although the majority of ribosomes, likely over 90% in bacterial cells^11^, are actively engaged in translation, a substantial fraction of mRNAs remains untranslated^12^. These observations imply that ribosomes are scarce relative to mRNAs and emphasize the need for their stringent allocation. Moreover, the proportion of actively translated mRNAs varies across cell types and physiological states^13^, which raises a critical question: whether and how RNA structural barriers prioritize and alter mRNA access to the finite ribosome pool.

Meiosis, a conserved eukaryotic developmental process, has been extensively studied in *Saccharomyces cerevisiae*. A cascade of transcription factors are critical for transcriptional regulation during meiosis progression^14^, nonetheless, transcriptional regulation is likely limited when chromosomes are undergoing movement and condensation, a defining feature of meiosis. Ribosome profiling studies revealed that translation is highly regulated during meiosis^15^ by mechanisms involving modulation of RNA-binding protein interactions^16^, upstream open reading frame usage (uORFs)^17^, and transcript isoform switching^18^. However, the mRNA structural landscape during meiosis remains uncharacterized, leaving the regulatory roles of RNA folding unexplored.

In this study, we applied dimethyl sulfate probing analyzed by mutational profiling (DMS-MaP) and RNA-seq to profile mRNA structure throughout meiosis in *S. cerevisiae*, covering seven meiotic stages spanning two nuclear divisions, and two control states of vegetative growth and pre-meiosis. The unique attributes of budding yeast, including reduced cellular heterogeneity, intron-sparse genome, and meiosis synchronization protocols^19^, enabled high-resolution determination of RNA structure. We found that transcripts upregulated during meiosis adopted more flexible structures than did the same transcripts during vegetative growth. RNA structural complexity and translation efficiency were more strongly correlated during meiosis than during vegetative growth, with complex structures reducing ribosome processivity in 5′ UTR and coding regions. Translation of structured mRNAs, including abundant transcripts like *CCW22*, was delayed to later stages of meiosis. We validated the role of RNA structural elements in governing *CCW22* translation timing using data-directed structure modeling and in-cell reporter systems.

We observed an oscillation in cytoplasmic RNA helicase levels from high to low to high as meiosis progressed, which paralleled a corresponding oscillation in the translational preference for mRNAs with distinct structural complexities. Overexpression of Ded1p helicase disrupted the translation timing of transcripts upregulated during meiosis in an RNA structure-dependent manner, causing meiosis progression to pause at metaphase II. In sum, our study indicates that intrinsic mRNA structures and oscillatory helicase levels coordinate translation timing and meiotic proteostasis in a comprehensive and cell-wide way. This mechanism emphasizes that RNA structure functions to control translation of individual messages, an increasingly well-validated mechanism, and also to integrate a translational program across hundreds of mRNAs. This pan-transcript mechanism is likely especially important during meiosis when gene regulation at the transcriptional level is compromised by chromosome condensation and movement.

## RESULTS

### High-resolution DMS-MaP Resolves mRNA Structures Across Yeast Meiosis

To generate comprehensive mRNA structure profiles of the yeast meiotic program, we used synchronous meiosis to achieve stage-specific resolution and sampled multiple time points for transcriptome probing. Meiosis was synchronized by inducing expression of the transcription factor Ndt80p with estrogen^19^ at pachytene, and samples were collected across seven meiotic stages, staged by spindle morphology imaging^20^ (Fig. S1A). We collected datasets at pachytene (T0), end of prophase I (T1), metaphase I (T2), anaphase I (T3), metaphase II (T4), anaphase II (T5), and meiotic exit (T6). In addition, we included vegetative growth (V) and pre-meiotic (P) samples as controls. All samples were analyzed using mRNA-seq and in-cell DMS-MaP.

DMS-MaP measures the chemical accessibility of RNA to DMS, a reagent that preferentially modifies unpaired adenosines and cytidines. We generated a comprehensive DMS-MaP dataset for the *S. cerevisiae* transcriptome across meiotic stages and in controls (Fig. 1A). The dataset is comparable in size (∼180 million reads per stage, Table S1) to large datasets obtained in studies of mammalian cells^8,21^. We quantified DMS reactivities at single-nucleotide resolution^22^. DMS treatment selectively modified adenosine and cytidine residues (Fig. S1B) and assessment of Pearson’s r value for DMS reactivities between replicates revealed that a read-depth threshold of 350 yielded reproducible structure modeling (Fig. S1C). The DMS-MaP data showed excellent agreement with structures of well-characterized RNAs like the 5S rRNA^23^ and *GLN1* mRNA^5^ (Fig. S1D).

**Figure 1.**
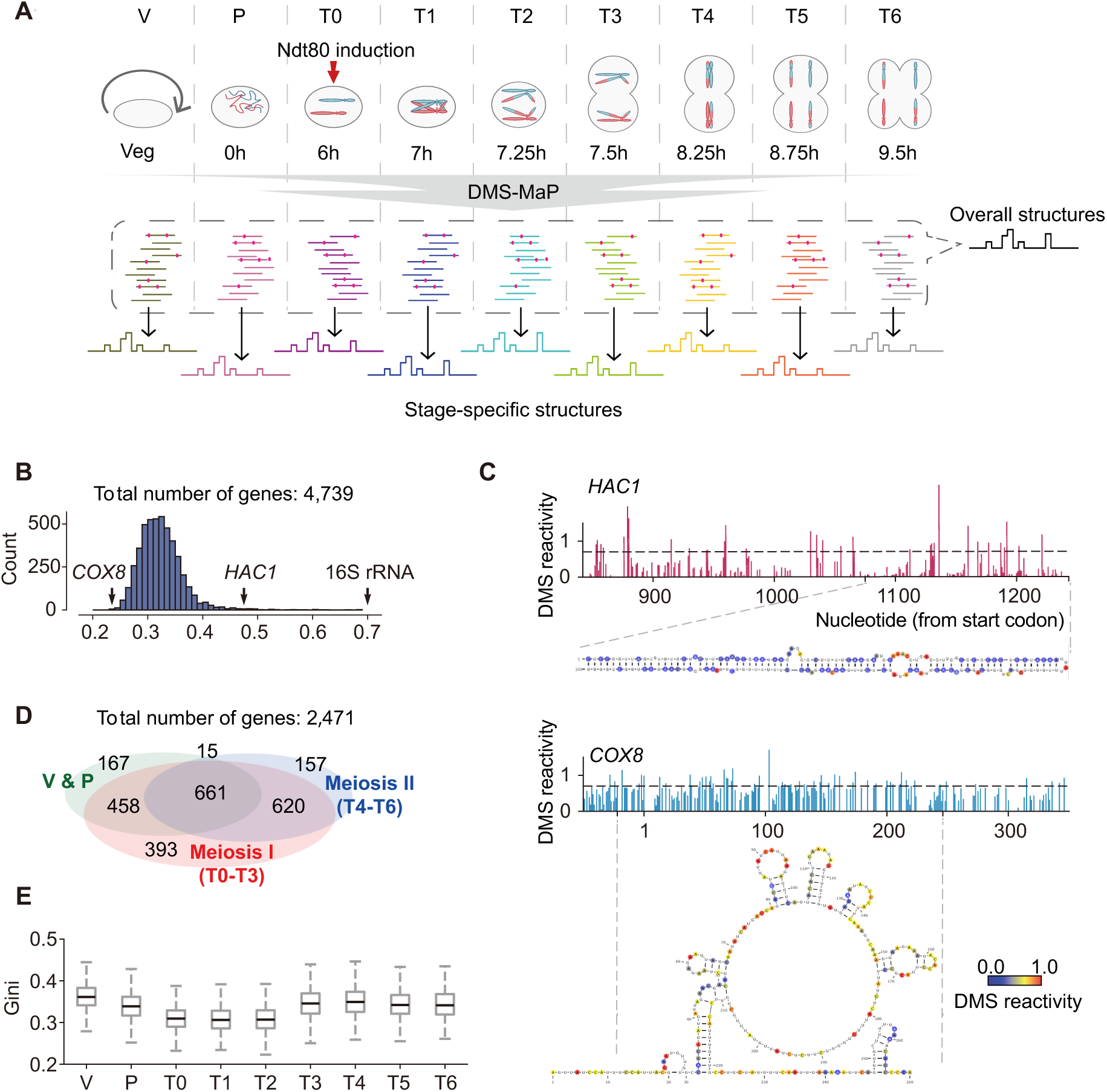
High-resolution mRNA structurome during yeast meiosis. (A) Experimental strategy to obtain mRNA structural maps during yeast meiosis. (B) Gini index distribution of mRNA structures from 4,739 genes. (C) DMS reactivities of a 400-nt region in *HAC1* and *COX8* mRNA and corresponding structure models. (D) Venn diagram of the numbers and relationships of stage-specific structures resolved during three periods: vegetative to pre-induction (V-P), early meiosis (T0-T3), and late meiosis (T4-T6). (E) Gini index distributions along the meiosis progression.

We performed quantitative structural analysis using two approaches. First, we pooled data across all stages to generate high-quality structures for 4,739 of 6,666 annotated yeast mRNAs (Table S2), with a median signal-to-noise ratio of 5.8 (Fig. S1E). For 3,399 mRNAs, more than 80% of adenosine and cytosine residues in open reading frames (ORFs) were detected (Fig. S1F). To quantify structural complexity, we calculated the Gini index, where higher values indicate more extensive DMS protection^5^. As expected, high Gini values (∼0.7) were observed for (highly structured) rRNAs (Fig. 1B). Structural complexity varied greatly among mRNA species (Fig. 1B). For example, *HAC1* mRNA, previously shown to form extensive intramolecular base-pairings^24,25^, had a high Gini index of 0.47 (Fig. 1C), whereas *COX8* mRNA, predicted to be largely single-stranded, had a Gini index of 0.24 (Fig. 1C). As a normalized measure of RNA structural complexity, the Gini index permits comparisons across samples^26^. For 2,471 mRNAs, structures could be modeled or quantified at more than one stage (Fig. 1D), and structures of 1,799 mRNAs were obtained at three or more stages (Table S3). We found that global Gini index initially decreased and then increased during progression across meiosis (Fig. 1E). Specifically, overall mRNA structure, as reported by mean Gini index, decreased significantly from vegetative growth to pachytene (V to T1), remained low until metaphase I (T2), and then increased substantially at anaphase I (T2-T3), remaining at this level until meiotic exit (T5).

### Transcripts Upregulated during Meiosis Adopt Simpler Folding Than Other mRNAs

Meiotic entry is accompanied by massive production of new mRNAs^14,27^, which we reasoned might lead to the global decrease in RNA structural complexity (Fig. 1E). Our mRNA-seq identified 1,078 transcripts that were upregulated during meiosis (TUMs) by at least eight-fold in abundance upon entry into meiosis. Transcripts that decreased relative to the vegetative state or changed by less than eight-fold were classified as non-TUMs (5,520 in total). TUMs fell into clear groups, which we categorized as early and late groups based on their peak abundance around meiosis I or meiosis II, respectively (Fig. 2A).

**Figure 2.**
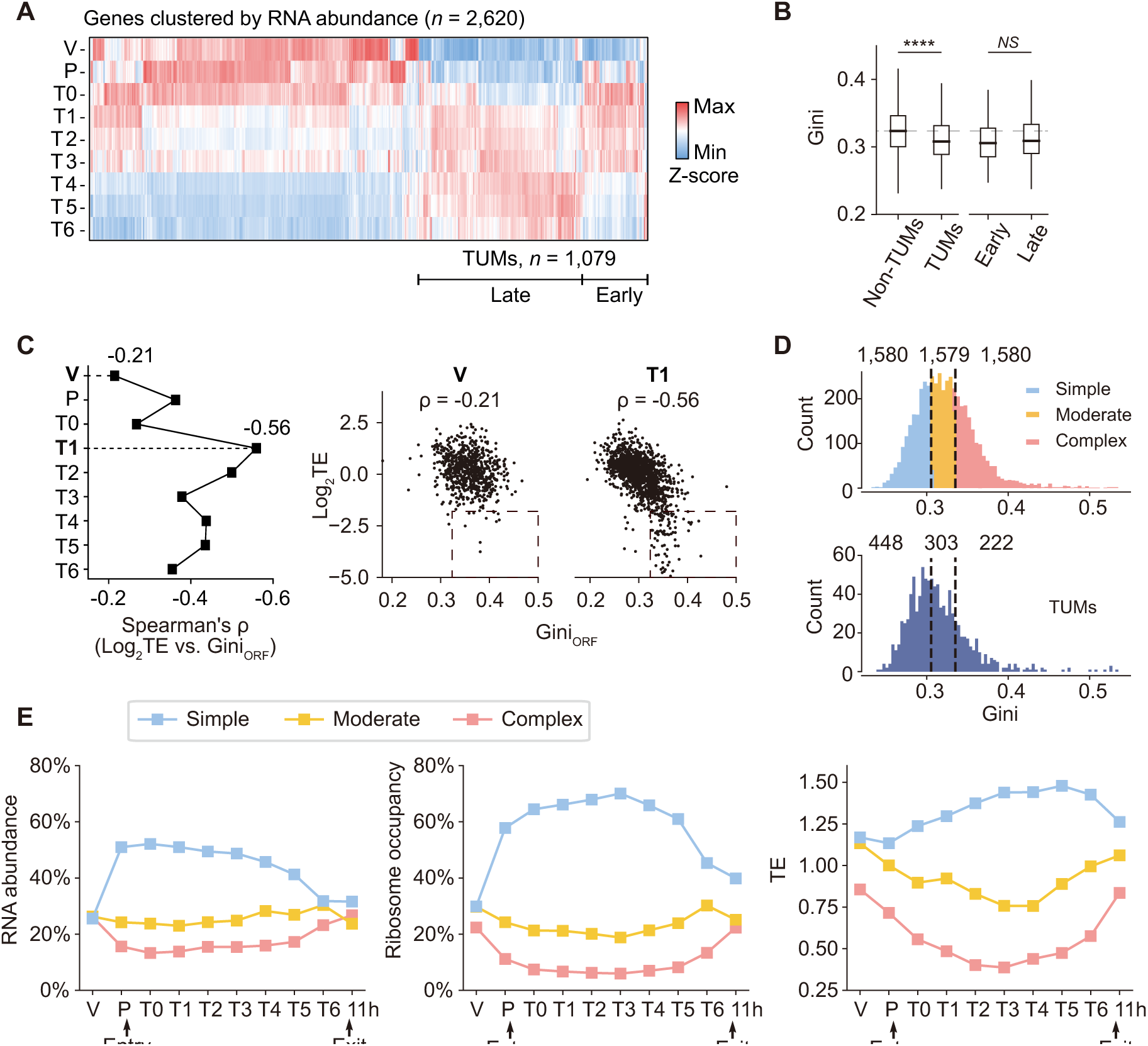
mRNAs with flexible folding are preferentially transcribed and translated during meiosis. (A) Hierarchical clustering of mRNAs (*n* = 2,620) with more than an 8-fold change (*P* < 0.05) in transcripts per million (TPM) during meiosis. (B) Distribution of Gini indices in non-TUM (*n* = 3,767), TUM (*n* = 937), Early TUM (*n* = 251), and Late TUM (*n* = 705) groups. The dashed line indicates the median Gini index for the non-TUM group. Statistical significance, non-TUM vs. TUM (\*\*\*\**P* ≤ 10^-23^) (C) Left: Spearman’s rank correlation coefficients between Gini indices and TEs across vegetative (V) to T6 stages. Right: Relationship between Gini indices and TEs at stage V (with the lowest correlation, n = 736) and stage T1 (with the highest correlation, n = 1,480). (D) Three equal-sized mRNA groups were defined based on RNA structural complexity: Simple, Moderate, and Complex. The number of TUMs in each group is indicated along with the corresponding cutoffs. (E) Proportional contributions of Simple, Moderate, and Complex mRNA groups to RNA abundance (left), ribosome occupancy (middle), and TE (right). RNA abundance was scaled to 100% to measure the proportion for each group. Similarly, ribosome footprints were normalized to indicate ribosome occupancy per group. The overall TE for each group was calculated by normalizing ribosome occupancy to mRNA abundance proportion.

We found that TUMs had significantly lower Gini indices than non-TUMs (*P* < 10^-23^, Fig. 2B), and no significant Gini index differences were noted between early and late subgroups of TUMs (*P* = 0.061, Fig. 2B). Thus, as cells enter meiosis, the downregulation of more structured non-TUMs and the upregulation of less structured TUMs collectively reduce global mRNA structural complexity. The subsequent downregulation of early-group TUMs in later meiotic stages may account for the recovery of global structural complexity (Fig. 1F).

The observed differences in structural complexity do not reflect simple differences in GC content. First, there are no significant differences in GC content within ORFs of TUMs and non-TUMs (*P* = 0.4, Fig. S2A). Second, we calculated Gini indices using only adenosine reactivities. The TUMs still had significantly lower indices than did non-TUMs, indicating that there was less base pairing of adenosines in TUMs than non-TUMs (Fig. S2B). The structural differences between TUMs and non-TUMs, revealed by the Gini index, are thus driven by intrinsic differences in mRNA folding. Given that RNA structures can impede translation and often require ATP-dependent unwinding, we sought to explore whether and how RNA structure in TUMs governs meiosis.

### mRNAs with Less Stable Structure are Preferentially Transcribed and Translated during Meiosis

A prior ribosome profiling study revealed widespread translation control throughout yeast meiosis^15^. Translation efficiencies (TEs) of certain mRNAs like *SPS1* varied up to 100 fold^15^. Although active translation has the potential to destabilize mRNA structures within coding regions, the extent of this effect in eukaryotes is unclear. We assessed the impact of translation itself on RNA structure in our system using matched time courses from ribosome profiling and our structurome datasets (Fig. S2C). We focused on RNA structural changes in a subset of transcripts with large TE fluctuations during meiosis^15^. For *ECM8*, *SPO20*, *SPS1*, *CLB3*, and *GIP1* mRNAs, which have TE variations ranging from 15 to 106-fold, DMS reactivity patterns were similar across meiosis (Fig. S2D). Further, analysis of ORF structural complexities (Gini_ORF_ index) of 390 transcripts with at least 4-fold TE changes showed no correlation between complexity and TE changes (r = −0.08; *P* = 0.16, Fig. S2E). These results suggest that, even in highly translated mRNAs, ORF structures are transiently disrupted but rapidly refold after ribosome passage. More specifically, internal RNA structure is an intrinsic feature of an mRNA, that can modulate translation, but is not fundamentally altered by translation.

We then investigated how intrinsic ORF structures influence translation by examining the genome-wide association between TE and Gini_ORF_. Complex, stable ORF structures are postulated to hinder translation elongation^28^. Consistent with this model, we observed negative correlations between Gini_ORF_ and TEs across all vegetative and meiotic stages (Spearman’s r: −0.21 ∼ −0.56, *P* < 10^-261^, Fig. 2C and Fig. S2F). The correlations were notably stronger during middle meiotic stages when TUMs are fully transcribed (T1-T2, Spearman’s r: −0.50 ∼ −0.56) than during vegetative growth (Spearman’s r: −0.21). The strongest correlation was detected at the end of prophase I (T1, Spearman’s r = −0.56, Fig. 2C), where mRNAs with high Gini indices trended to have very low TEs (Fig. 2C). Correspondingly, the correlation between RNA abundance and ribosome profiling weakened significantly as meiosis progressed (Fig. S2G), further supporting our hypothesis that RNA structure acts as a dominant regulator of translation during meiosis.

To validate the association between RNA structure and TE, we stratified genes into three equal-sized groups based on RNA structural complexity: Simple, Moderate, and Complex (Fig. 2D). There were twice as many TUMs in the Simple group (n = 448) as in the Complex group (n = 222) (Fig. 2D). Transcripts in the Simple group had significantly higher TEs than those in the Complex group across all stages, with the difference being more pronounced during meiosis than during vegetative growth. Specifically, the average TE of the Simple group was approximately 1.4-fold higher than that of the Complex group during vegetative growth, and this gap widened to over 2.0-fold during meiosis (Fig. S2H). Moreover, TUMs showed relatively low TEs during vegetative growth but experienced elevated TEs during meiosis, particularly at the early stages (T2 and T3, Fig. S2I). These results indicate that the general inhibitory effect of RNA structure on translation is specifically amplified during meiosis and that the meiotic cellular environment, which favors translation of unstructured RNAs, is especially permissive for TUM translation.

These results thus indicate that mRNAs with less structured folding are both preferentially transcribed (Fig. 2B) and translated (Fig. 2C) during meiosis. To quantify this preference, we next scaled RNA abundance and ribosome footprint signals to 100% at each time point and calculated the proportional contributions of transcripts stratified by structural complexity. This analysis revealed strong similarities, or potential coupling, among transcriptome composition, ribosome occupancy, and RNA structure (Fig. 2E). During vegetative growth, mRNAs with different structural complexities contributed similarly to the transcriptome and translatome. After entry into meiosis, however, transcripts with flexible folding are more highly expressed and become preferentially engaged by ribosomes, before returning to a more balanced distribution at meiotic exit (Fig. 2E). When we normalized the ribosome occupancy by mRNA abundance, a similar pattern in TE across meiosis was observed (Fig. 2E). These findings emphasize that structurally flexible TUMs are preferentially translated during middle meiosis.

The Simple and Complex groups showed structural differences across 5′ UTRs, ORFs, and 3′ UTRs, with the Simple group displaying consistently lower DMS reactivity (Fig. S3A). To assess how ORF structures contribute to the distinct translation patterns of the two groups, we calculated ribosome flux from the 5′ to the 3′ end in 50-nucleotide windows (Fig. S3B), and defined the most structured window (window 0) by the maximal Gini index (Fig. S3C). During vegetative growth, ribosome flux decreased similarly in both groups across the most structured window (Fig. S3D). In contrast, during meiosis, the Complex group showed a 10% greater reduction in ribosome flux than the Simple group (Fig. S3D), indicating that intricate local structures impose stronger barriers to ribosome progression during meiosis. Given that translation from uORFs in yeast occurs predominantly during meiosis^15^, we further examined whether 5′ UTR structure influences ribosome movement during meiosis. Ribosome footprinting revealed that the frequency of uORF translation in the Complex group was twice that in the Simple group (Fig. S3E), suggesting that 5′ UTR structures impede ribosome processivity and could promote the use of alternative start codon in 5′ UTR. Together, these results indicate that RNA structure shapes the divergent translation patterns of the Simple and Complex mRNA groups by modulating ribosome processivity in 5′ UTR and ORF regions.

### mRNA Structure Contributes to Stage-Specific Translation

A central question in meiosis is how pre-transcribed mRNAs are selectively translated at different meiotic stages. Although TUMs generally have simpler structures than non-TUMs, their structural complexity varies considerably (Fig. 3A). To explore whether RNA structures influence the translation timing of TUMs, we classified them into three equal groups based on Gini indices and compared their ribosome occupancies at each stage. The Simple group peaked in ribosome occupancy around anaphase I (T3: 40%), whereas the Complex group showed a delayed surge only during meiotic exit (T6: 20.5%). The Moderate group had a pattern that fell between these two extremes (Fig. 3A). This association was more pronounced for structural complexity in 5ʹ UTR and ORF regions than in 3ʹ UTR regions (Fig. S4A). Gene Ontology analysis revealed stage-specific functional associations: Genes involved in early or middle-stage events (synapsis, recombination, chromosome segregation) were predominantly in the Simple and Moderate groups, whereas genes in the Complex group were primarily associated with cell wall activities (Fig. S4B and Table S4), functions critical for gamete maturation after meiosis II. Moreover, genes known to remain untranslated until the end of meiosis II^29^, such as *SPS1*, *SPS4*, and *SSP2*, were significantly enriched in the Complex group (*P* < 10^-7^, Table S5). These findings support the hypothesis that RNA structure dictates the temporal translation of TUMs to ensure stage-specific functions.

**Figure 3.**
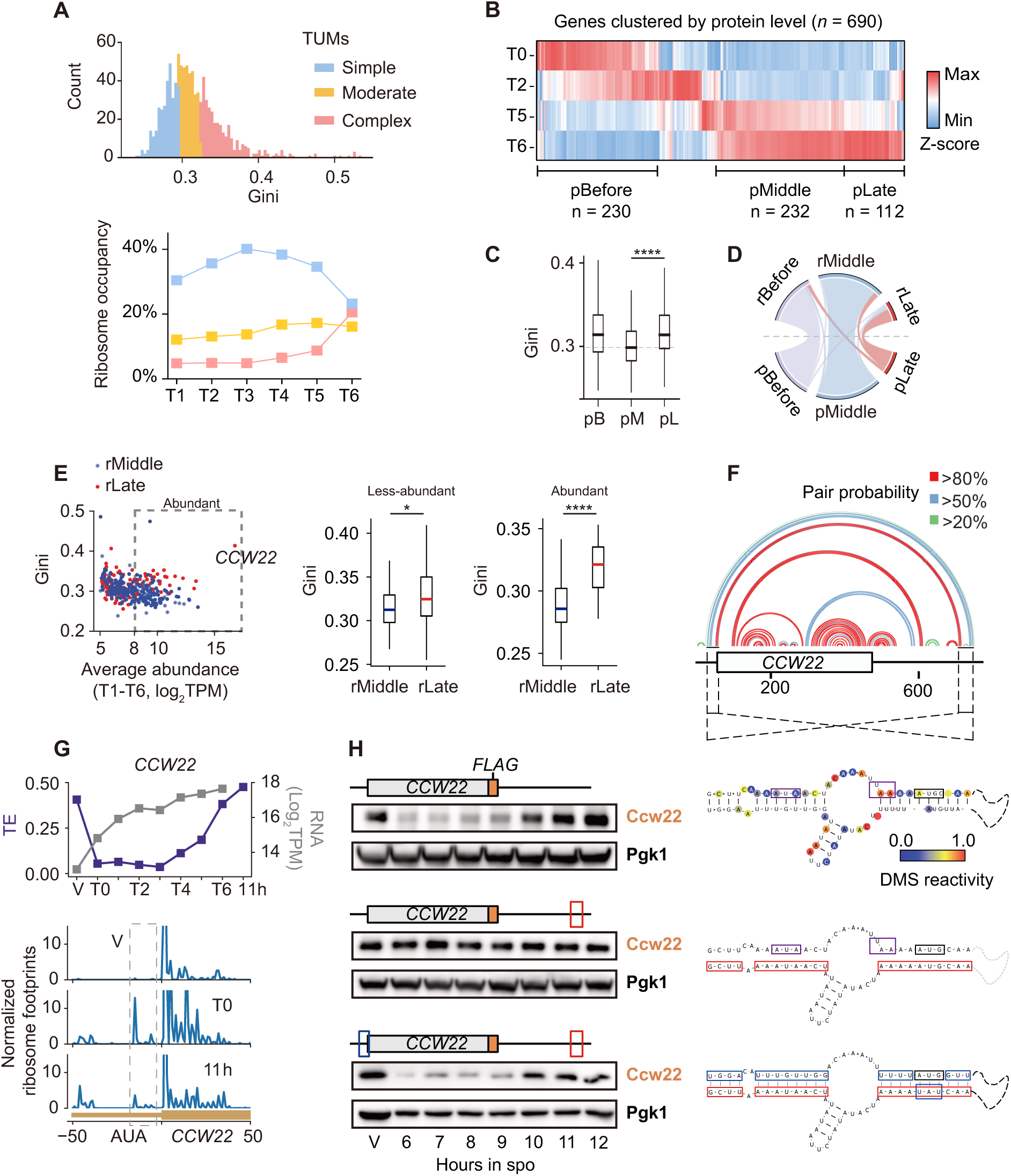
mRNA structure regulates translation timing. (A) Distribution of Gini indices for TUMs (upper) and dynamic ribosome occupancy (lower) for mRNAs grouped based on structural complexity: Simple (*n* = 324), Moderate (*n* = 325), and Complex (*n* = 324). (B) Hierarchical clustering of protein abundances for genes exhibiting more than 1.2-fold changes (*n* = 690, *P* < 0.05). (C) Comparison of Gini indices among three protein groups: before meiosis (pBefore, *n* = 192), during mid-meiosis (pMiddle, *n* = 223), and near meiotic exit (pLate, *n* = 104). (D) Overlap analysis between genes in ribosome profiling and protein clusters: Before, *P* < 10^-38^; Middle, *P* < 10^-50^; Late, *P* < 10^-18^. (E) Left: Relationship between Gini indices and average mRNA abundance (log_2_TPM) during meiotic stages T1-T6 for transcripts with different translation timings (rMiddle, rLate). Abundant transcripts were defined as those with TPM > 8. Right: Comparison of Gini indices between abundant and less-abundant transcripts within the rMiddle and rLate groups. (F) Data-directed structure model of *CCW22* mRNA shown as an arc plot with arcs colored based on base-pairing probability. (G) *CCW22* mRNA abundance and TE changes during meiosis (upper), and ribosome footprints on *CCW22* uORF at time of high TE (V and 11 h) and low TE (T0) (lower). The ribosome footprint reads were normalized as in described in the Fig. 2H legend. (H) Western blot analysis of Ccw22 protein expressed from wild-type (top), structure-disrupted (middle), and structure-restored (bottom) vectors. The red boxes indicate mutated nucleotides. The blue boxes indicate nucleotides mutated to restore base pairing. The purple boxes indicate the start codon and stop codon of a *CCW22* uORF.

To evaluate the role of RNA structure in the expression timing of individual proteins during meiosis, we conducted quantitative proteomics across four meiotic stages: pachytene (T0), metaphase I (T2), anaphase II (T5), and meiotic exit (T6). Among 3,989 quantified proteins, 690 changed significantly (>1.2-fold, *P* < 0.05, Table S2). These proteins were clustered into three groups based whether abundance peaked before meiosis, during mid-meiosis, or near meiotic exit (Fig. 3B). Proteins that peaked in mid-meiosis had significantly simpler mRNA structures than those that peaked before meiosis or near meiotic exit (Fig. 3C), which supported a connection between mRNA structure and protein expression dynamics during meiosis.

We next classified genes with greater than 8-fold changes (*n* = 1,666) in ribosome footprints across meiosis into three groups according to their translation profiles (Fig. S4C). Genes grouped by translation timing significantly overlapped with those classified by protein expression dynamics (before, *P* < 10^-38^; mid-meiosis, *P* < 10^-50^; near meiotic exit, *P* < 10^18^; Fig. 3D), indicating that translational timing is a major determinant of protein expression dynamics. Consistent with the trend observed for protein clusters, transcripts with peak translation near meiotic exit (n = 217), including Ndt80 targets known for delayed translation such as *CLB3*^19^ and *SPS1*^29^, had higher Gini indices than transcripts translated during mid-meiosis (*P* < 10^-11^, Fig. S4D), and were enriched in the Complex group (*P* < 10^-7^, Table S6). Regional analysis revealed that structural differences were concentrated in the 5ʹ UTRs and ORFs (*P* = 0.016 and *P* < 10^-9^, respectively), but not in 3ʹ UTRs (Fig. S4E and Table S7). RNA-seq profiles revealed that these genes were transcribed from pachytene to metaphase II (Fig. S4F), highlighting varying degrees of translation delays. Together, these results suggest that increased structural complexity within the 5′ UTR and ORF regions contribute to delayed translation at late stages.

### Validation of Long-Range RNA Structure in Regulating *CCW22* Translation Timing

Given that highly abundant mRNAs place substantial demands on ribosome resources, we examined whether their translation is more strongly constrained by RNA structure than mRNAs of lower abundance (Fig. 3E). Intriguingly, for abundant mRNAs, translation timing was more significantly associated with Gini indices (Fig. 3E). For example, *CCW22*, which encodes a cell wall protein, is the most abundant mRNA during meiosis. It is translated near meiotic exit and has an exceptionally high Gini of 0.414 (Fig. 3E). These observations indicate that the translation timing of abundant mRNAs is more tightly regulated through RNA structure.

DMS-MaP–guided structural modeling showed that the *CCW22* mRNA possesses a highly compact ORF structure and a long-range interaction between the 5′ UTR and the 3′ UTR (Fig. 3F). Despite continuous accumulation and high abundance during meiosis, *CCW22* mRNA has a high TE only during vegetative growth and at late meiotic stages (Fig. 3G). Ribosome footprint analysis revealed apparent ribosome stalling at a near-cognate start codon (*AUA*) at position −17 in its 5′ UTR, specifically during low TE stages (Fig. 3G). This ribosome stalling site resides within the base-paired region resulting from the 5′ UTR*–*3′ UTR interaction (Fig. 3H). Together, these observations suggest that the long-range 5′ UTR*–*3′ UTR structure of *CCW22* may regulate its translation timing.

We created reporter constructs for expression of the wild-type *CCW22* mRNA and mutant mRNAs. First, deletion of either the entire 5′ UTR or the 3′ UTR led to earlier protein expression compared to the wild-type construct (Fig. S5A). Second, more precise mutagenesis of 3′ UTR nucleotides involved in base-pairing with the 5′ UTR similarly accelerated Ccw22 protein expression (Fig. 3H). Restoring the interactions through compensatory mutations in the 5′ UTR restored expression timing to that of the wild-type construct (Fig. 3H). All constructs produced mRNAs that accumulated rapidly during early meiosis (Fig. S5B), ruling out transcriptional effects. These results show that the long-range structure of *CCW22* mRNA governs its translation timing.

Consistent with its extreme abundance, ribosome occupancy for *CCW22* increased from 0.05% during pachytene to 4.2% at meiotic exit (Fig. S5C). The pronounced engagement of ribosomes by a single gene suggests that structural elements within *CCW22* could influence global translation events. Supporting this model, a yeast strain expressing a *CCW22* variant unable to form base pairs between the 5′ UTR and the 3′ UTR had a significant delay in meiosis progression compared to the wild-type strain (Fig. S5D). This finding indicates that failure to restrain translation of highly abundant transcripts can exert cell-wide effects, presumably through competitive access to a finite ribosome pool, underscoring the role of RNA structure in ensuring proper meiotic progression.

### Helicase Expression Is Correlated with Translation Timing during Meiosis

We hypothesized that changes in allocation of ribosomes may arise from fluctuations in cellular sensitivity to mRNA structure. RNA helicases are the main factors responsible for unwinding RNA structures *in vivo*^30^. In mammals, helicases such as YTHDC2, DHX36, and DDX25 are important at various stages of meiosis and spermatogenesis^31–33^, suggesting that helicase abundance and activity may be precisely and dynamically modulated across developmental stages.

We analyzed the expression of 14 cytoplasmic RNA helicases and 2 helicase-regulating accessory proteins. Because our dataset does not include proteomic measurements from yeast cells during vegetative growth or from mature gametes, we incorporated published protein abundance data from the SK1 yeast strain collected during natural meiosis^18^ (Table S8). Co-clustering of helicase expression at the translation and protein levels revealed two distinct patterns (Fig. 4A). Levels of helicases such as Ded1 continuously declined from vegetative growth through gamete maturation, whereas those in the other group, including Dbp1, were detected at low levels in early meiosis and increased in later stages. Collectively, the protein expression of these helicases oscillates in a high-low-high pattern over the course of meiosis.

**Figure 4.**
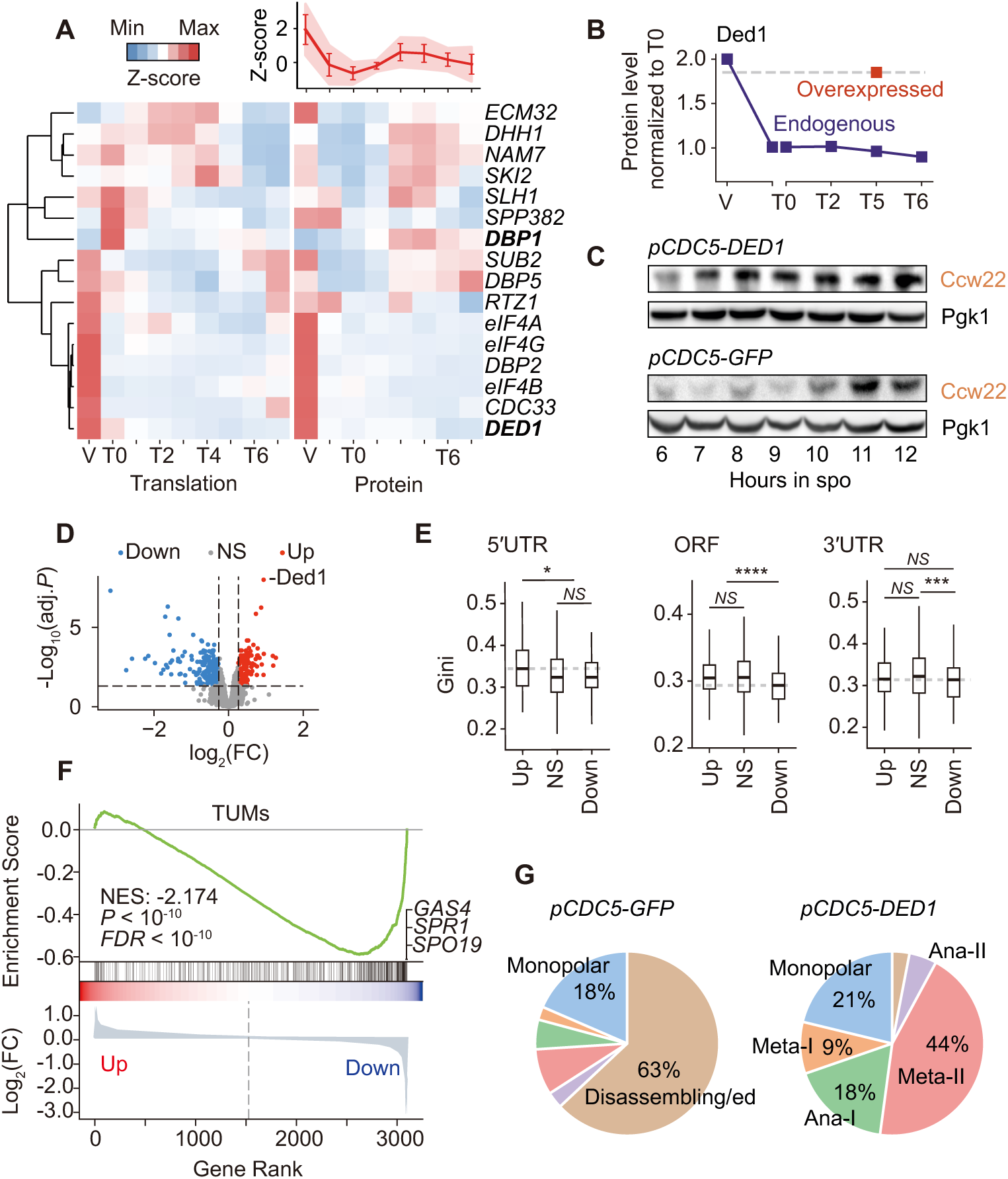
Helicase expression governs meiotic protein expression. (A) Co-hierarchical clustering of 14 cytoplasmic RNA helicases and 2 accessory proteins during meiosis. Ribosome profiling data (left) were from synchronized meiosis, and proteomic data (right) were from the strain undergoing natural meiosis^18^. Mean protein abundance (Z-scored) of all helicases is shown in the upper right. (B) Abundances of endogenous Ded1 protein (blue squares) and Ded1 protein overexpressed under the control of the *CDC5* promoter (red squares) at T5 stage. (C) Western blot analysis of Ccw22 expression in strains with Ded1 or GFP overexpression, both under the control of the *CDC5* promoter. (D) Volcano plot of genes differentially expressed in cells that overexpress Ded1 with log_2_ fold change > 1.2 and P < 0.05. Upregulated, n = 149; downregulated, n = 194; not significant, n = 2,761. (E) Comparison of Gini indices across 5′ UTR, ORF, and 3′ UTR regions of mRNAs differentially expressed under Ded1 overexpression. (F) Gene Set Enrichment Analysis of proteins expressed by TUMs that that are up- or downregulated by Ded1 overexpression. (G) Percentages of cells in indicated meiotic stages at 12 hours post-induction in yeast strains overexpressing GFP or Ded1.

Independent support for a role for the interplay of helicase expression and RNA structure during meiosis comes from our analysis of the correlation of mRNA structure and TE. Specifically, mRNAs with low Gini indices were preferentially translated during early meiotic stages, when RNA helicase expression is suppressed, whereas structurally complex transcripts required higher helicase levels for efficient translation and therefore reached peak translation during late meiosis when RNA helicase expression partially recovers (Fig. 2E). Notably, protein expression from the highly structured *CCW22* transcript closely mirrored the oscillation of RNA helicase expression, with elevated levels only during vegetative growth and late meiosis (Fig. 3H).

### Overexpression of Ded1 Helicase Disrupts Protein Levels during Meiosis

To test our model that helicase activity directs protein expression, we altered the RNA helicase level in early meiosis by overexpressing Ded1, a helicase that resolves 5′ UTR structures^34^. In unperturbed cells, ribosome profiling and proteomic data revealed Ded1 levels decrease by 50% from vegetative growth (V) to pachytene (T0) and remain low through the end of meiosis (Fig. 4B and Fig. S6A). To elevate Ded1p levels, we placed *DED1* under the control of the *CDC5* promoter, which drives robust expression upon meiotic induction (Fig. S6B). Under control of the *CDC5* promoter, Ded1p rose to a near-vegetative level by anaphase II (Fig. 4B). In these cells overexpressing Ded1p, Ccw22 was expressed prematurely in early meiosis I (Fig. 4C). This result is exactly consistent with a role for Ded1p in overcoming structural barriers within the *CCW22* 5′ UTR (Fig. 3F).

Ded1p overexpression should further cause misassociation of ribosomes in an RNA structure-dependent manner, leading to inappropriate protein expression during meiosis. To test this prediction, we conducted differential protein expression analysis comparing wild-type and Ded1-overexpressing strains. This comparison identified 149 genes that were upregulated and 194 genes that were downregulated by at least 1.2-fold upon Ded1 overexpression (*P* < 0.05; Fig. 4D and Table S9). Critically, upregulated genes had more complex mRNA structures than downregulated ones (*P* < 10^-5^; Fig. 4D). Region-specific comparisons showed that upregulated mRNAs had more highly structured 5′ UTRs than did downregulated mRNAs (*P* = 0.014, Fig. S6C), consistent with the function of Ded1 in unwinding 5′ UTR secondary structures. Conversely, downregulated mRNAs had simpler 5′ UTR and ORF structures (*P* < 10^-5^, Fig. S6C), suggesting that their suppression is a consequence of ribosome reallocation toward more structured mRNAs. These findings support a model in which Ded1 overexpression globally shifts the translational landscape toward mRNAs with more complex structures.

Because TUMs generally adopt flexible structures and are preferentially translated during low-helicase stages, we evaluated how elevated Ded1p levels influence TUM expression and meiotic progression. Gene Set Enrichment Analysis revealed a decrease in abundances of TUM-encoded proteins upon Ded1 overexpression (Fig. 4E), with the strongest reductions observed for the most structurally flexible TUMs, including *SPO19*, *SPR1*, and *GAS4* (Gini indices of 0.28, 0.27, 0.27, respectively). Moreover, Ded1 overexpression severely disrupted the meiotic cell cycle, frequently causing arrest at or before metaphase II, as revealed by immunofluorescence staging at 12 hours postinduction (Fig. 4F). Together, these findings suggest that helicase expression levels coordinate ribosome allocation among structurally distinct mRNAs, thereby shaping stagespecific gene expression that is crucial for faithful execution of meiotic program (Fig. 5).

**Figure 5.**
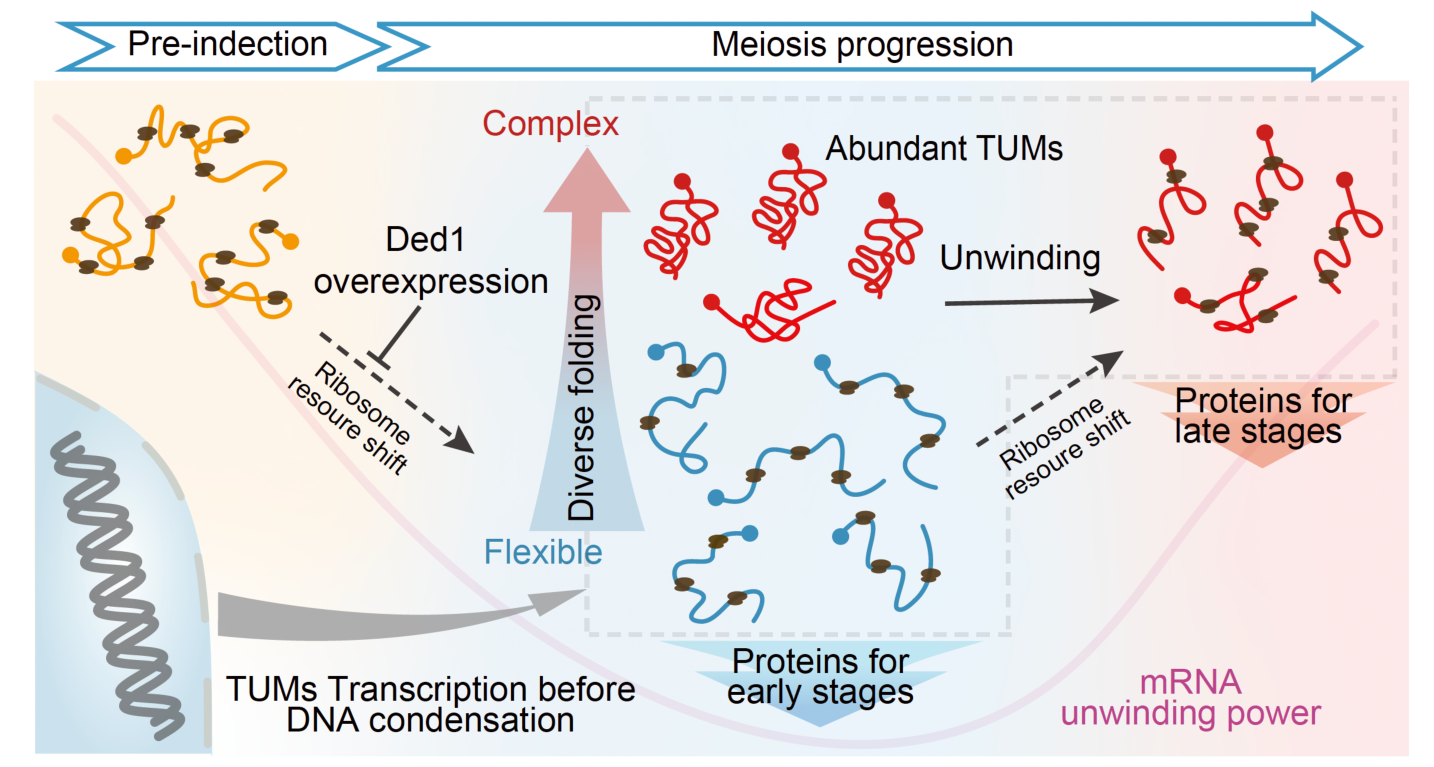
RNA structure and mRNA unwinding power orchestrate stage-specific translation. Model for how diverse RNA structures and oscillatory mRNA unwinding power synergistically orchestrate the dynamic allocation of ribosome resource. These combined features enable selective translation and precise protein synthesis during meiosis.

## Discussion

High-throughput RNA structure analyses in eukaryotes require deep sequencing to detect low-abundance transcripts. This challenge is even greater when applied to dynamic processes that demand time-resolved resolution. Most studies to date have focused on a few cell states or individual RNA elements, leaving a gap in our understanding of how RNA structure coordinates cellular processes across time. Here, we generated a high-resolution mRNA structural map across nine stages of synchronized yeast meiosis, covering approximately 70% of annotated mRNAs. Integrating this dataset with parallel multi-omic measurements revealed that RNA folding regulates selective gene expression across meiotic stages to ensure the accurate execution of meiosis (Fig. 5).

Through combined analyses of RNA structure with mRNA abundance, translation efficiency, translation timing, and protein expression dynamics, we uncovered four key correlations. First, transcripts upregulated during meiosis (TUMs) tend to adopt more flexible structures. Second, mRNAs with flexible structures are translated more efficiently during meiosis, whereas translation of highly structured mRNAs is delayed until late meiosis. Third, complex 5′ UTR and ORF structures correlate with later translation timing. Finally, cytoplasmic RNA helicase expression smooths out the preference for translating RNA with lower levels of structure, enabling highly structured mRNAs to be translated efficiently. Together, these correlations support a model in which RNA structure and RNA helicases jointly regulate stage-specific translation: When cells enter meiosis, a large cohort of mRNAs (TUMs) is transcriptionally induced. Although these mRNAs are generally unstructured, they exhibit considerable diversity in structural complexity. During early meiosis, the global suppression of helicases restricts the translation of highly structured mRNAs while biasing ribosomes toward mRNAs with flexible structures, including most TUMs. This selective translation of newly transcribed TUMs facilitates rapid translational reprogramming into a meiotic-specific mode. During late meiosis II, specific helicases are induced, which increases translation of highly structured mRNAs. We propose that this RNA structure-based regulation is particularly adaptive for the cell when chromosome compaction limits new transcription during meiotic divisions.

We directly validated this model from the perspectives of both RNA structure and RNA helicases. For the most abundant transcript expressed during meiosis, *CCW22*, we confirmed that a specific RNA structure, a long-range 5′ UTR–3′ UTR interaction controls its protein expression timing. Disruption of this RNA structure led to ribosome monopolization at inappropriate stages and delayed meiotic progression. Restoring the RNA structure rescued normal Ccw22p expression timing. Strikingly, overexpression of the Ded1 helicase disrupted gene expression in an RNA structure-dependent manner and led to complete meiotic arrest. Proteomic profiling revealed increased abundance of several helicases implicated in mRNA translation during late meiosis. For example, Dbp1p, a “low performance” paralog of Ded1^35^, may unwind structured 5′ UTRs, whereas Dhh1 may facilitate translation of structured ORFs^36^. The roles of these helicases during late meiosis remain compelling opportunities for future investigation.

Our study supports a systemic framework in which mRNAs compete for a finite pool of ribosomes governed by their intrinsic folding, while the complement of expressed helicases – or cellular unwinding power – modulates the extent to which RNA structure influences translation. This zero-sum framework provides a unified explanation for how RNA structure coordinates translation across multiple transcripts. For instance, our model accounts for the observation that Ded1 inactivation suppresses translation of mRNAs with structured 5′ UTRs while simultaneously enhancing translation of mRNAs with simple 5′ UTRs^37^. By contrast, focusing on individual transcripts makes it difficult to explain enhanced translation of specific mRNAs under helicase inactivation. Our model attributes this dual effect to a transcriptome-wide reallocation of ribosomes from highly structured to more flexible mRNAs, as directly supported by our Ded1 overexpression experiments.

The RNA structure-helicase interplay model applies broadly to biological processes in which functionally related mRNAs share similar structures and RNA helicases undergo significant changes. Both conditions have been widely documented in other biological systems. Genes that participate in environmental response processes encode mRNAs with more complex structures, including double-stranded structures downstream of upstream AUGs in immune response genes^10^ and structured regions that occur shortly after start codons in autophagy genes^36^. These structural features provide the basis for large-scale coordinated expression regulation based on RNA structure. Furthermore, fluctuations in helicase expression have been implicated in other physiological and pathological contexts, including innate immune responses, neurodegenerative diseases, and multiple cancers^38^. Therefore, helicase-mediated ribosome reallocation can serve as a general post-transcriptional mechanism for coordinating gene expression across diverse biological processes.

## Data availability

The raw sequencing data are available at the NCBI BioProject under accession number PRJNA1174166. Scripts used for sequencing data processing and analysis are accessible at the GitHub repository: https://github.com/HoNg621/StructuromeYeastMeiosis. The mass spectrometry raw data have been deposited in the ProteomeXchange Consortium member repository with accession code PXD056964. Further inquiries regarding resources and reagents should be directed to the lead contact, H.W.

## Acknowledgments

This work was supported by the National Key Research and Development Program of China (2016YFA0500900 to Y.B.), start-up funding from ShanghaiTech University (to Y.B.), and the US National Institutes of Health (R35 GM122532 to K.M.W.). We thank the Molecular and Cell Biology Core Facility (ShanghaiTech University), the Molecular Imaging Core Facility (ShanghaiTech University), and the Multi-Omics Core Facility (ShanghaiTech University) for technical support. We are indebted to Yang Yang (ShanghaiTech University) for critical comments. We thank Wei Li (Guangzhou Women and Children’s Medical Center) for sharing yeast strains, and Jilong Liu (ShanghaiTech University) for sharing plasmids.

## Author contributions

Y.B. conceived of and supervised the study. H.W. and Y.B. designed the experiments with assistance from C.Z. H.W. and Y.B. designed the analyses. H.W. and C.Z. performed the experiments with assistance from S.D. and Y.X. H.W. performed the analyses. H.W., Y.B., and K.M.W. wrote and revised the manuscript with contributions from S.F.

## METHODS

### Synchronous Meiosis Conditions

The engineered yeast strain A14201 (SK1 background), commonly used in meiotic studies, was cultured to saturation in YPD medium (1% yeast extract, 2% peptone, 2% glucose). This culture was then diluted to an optical density at 600 nm (OD_600_) of 0.3 in 1% yeast extract, 2% peptone, 1% potassium acetate, pH 7.0 (YPA medium) and grown overnight. Cells were washed twice and resuspended in 0.5% potassium acetate, pH 7.0, 0.02% raffinose (SPO medium) to an OD_600_ of 1.9. At 6 hours after transfer to SPO, 1 µM β-estradiol was added to induce expression of the meiotic transcription factor Ndt80.

### Meiotic Staging

Tubulin immunofluorescence was performed to monitor spindle morphology as previously described^20^ with minor modifications. Cells were fixed with 3.7% formaldehyde for 20 minutes at room temperature, washed twice with 1x PBS, and resuspended in 100 μl of 1 M sorbitol in PBS. Cell walls were digested with 0.2 U/μl lyticase (Yeasen, 10403ES81) for 20 minutes at 30 °C for vegetative cells and 30 minutes at 30 °C for sporulating cells. The cells were then adhered to polylysine-treated slides and immersed in methanol at −20 °C for 6 minutes, followed by 30 seconds in acetone at room temperature. After air drying, cells were incubated with 5% bovine serum albumin, and the yeast spindle was visualized using mouse anti-alpha-tubulin antibody (Sigma, T5168, 1:1,000) as the primary antibody and Cy5-conjugated goat anti-mouse IgG (Jackson ImmunoResearch Laboratories, 115175-146, 1:200) as the secondary antibody with DAPI staining to identify the nuclear region.

### DMS-MaP Library Generation

For *in vivo* modification, 4 ml of yeast culture (OD_600_ of 0.8) were treated with 200 μl DMS for 4 minutes at 30 °C. Control samples were treated with an equal amount of DMSO. The reaction was quenched with 4 ml of ice-cold 30% beta-mercaptoethanol, and cells were quickly pelleted and washed with 2 ml ice-cold 30% beta-mercaptoethanol. Total RNA was isolated using a modified hot phenol method suitable for meiotic cells^39^ and treated with RNase-free DNase I (Promega, M6101) to remove genomic DNA. Poly(A) mRNA was enriched from 5 µg of total RNA using a Poly(A) mRNA Magnetic Isolation Module (NEB, E7490).

Reverse transcription and second-strand synthesis for mutational profiling were adapted from previous methods^21,40^, with modifications to preserve strand information. The first strand was synthesized using SuperScript II (Invitrogen, 18064014) with MaP buffer (50 mM Tris, pH 8.0, 75 mM KCl, 6 mM MnCl_2_, 10 mM DTT, 0.5 mM dNTPs) and strand-specific reagent (NEB, E7765). Following G-25 microspin column desalting (GE Healthcare, 27532501), the cDNA was compatible with the second-strand synthesis module in the directional RNA library prep kit (NEB, E7765), which included dUTP incorporation. After purification using SPRIselect beads (NEB, E7765), the double-stranded DNA was enzymatically fragmented using the DNA fragmentation module (NEB, E7810). Most fragments were roughly 400 bp in length. Further library generation procedures, including adaptor ligation, USER enzyme digestion, size selection (> 200bp, 1:1 ratio of beads to sample), and PCR amplification, were performed as per the manufacturer’s instructions (NEB, E7765).

### Tandem Mass Tag-based LC-MS/MS

For meiotic samples, yeast cultures (2 ml) were spun down and resuspended in 0.7 ml lysis buffer (20 mM HEPES, pH 7.5, 140 mM KCl, 2 mM EDTA, 1% Triton X-100, 1 × protease inhibitor (APExBIO, K1010)). Cells were then lysed by rigorous vortexing at 4 °C with acid-washed glass beads (0.4-mm diameter). Cell debris was removed by centrifugation. The crude proteins were precipitated by adding 10 volumes of cold acetone to the supernatant. After overnight incubation at −20 °C, the proteins were pelleted by centrifugation. After air drying, the pellet was dissolved in 100 µl 8 M urea in 50 mM NH_4_HCO_3_.

To disrupt disulfide bonds, dithiothreitol was added to a final concentration of 5 mM, and samples were incubated for 1 hour at 37 °C in a thermal mixer. Protein concentration was determined using a BCA assay (Yeasen, 20201ES76). Subsequently, 20 mg total protein was then alkylated with 10 mM iodoacetamide for 45 minutes at room temperature. Alkylated proteins were diluted 1:8 with 50 mM NaHCO_3_ and digested using trypsin protease (Life/Invitrogen, 90057) for 16 hours with an enzyme-to-substrate ratio of 1:50. The digestion was stopped by the addition of trifluoroacetic acid to a final concentration of 0.4%. The peptides were desalted over a C18 column and eluted in 70% acetonitrile, 0.1% formic acid. After evaporation to dryness in a vacuum concentrator, the sample was dissolved in 50 µl of 50 mM tetraethylammonium bromide.

Peptides were then labeled with 0.2 mg TMT10plex reagent (Thermo Scientific, 90309). The labeling reaction was stopped by adding 1 µl of 5% hydroxylamine. After desalting over a C18 column, peptides were eluted in 70% acetonitrile, 0.1% formic acid. After drying, the sample was dissolved in 500 µl of 50 mM tetraethylammonium bromide. Reverse-phase chromatography was used to reduce peptide complexity; 36 fractions were eluted with gradient elution buffer (increasing concentration of acetonitrile, 0.1% formic acid) and combined into eight fractions (1-2,10,18,26; 3,11,19,27; 4,12,20,28; 5,13,21,29; 6,14,22,30; 7,15,23,31; 8,16,24,32; 9,17,25,33-36). After evaporation to dryness, liquid chromatography-tandem mass spectrometry (LC-MS/MS) analyses were performed on a Q-Exactive HF-X.

### Western Blotting and RT-qPCR

Protein samples from yeast cultures were prepared and resolved via SDS-PAGE, followed by transfer to polyvinylidene difluoride membranes. Pgk1 and Ccw22-FLAG were detected using primary antibodies mouse anti-PGK1 (Abcam, ab113687, 1:10,000) and mouse antiFLAG (Sigma, F1804-50UG, 1:2,500), respectively. Peroxidase-conjugated goat antimouse IgG secondary antibody (Jackson ImmunoResearch Laboratories, 115-035-146, 1:10,000) and ECL luminescent solution (Share-Bio, SB-WB012L) were used for blot detection. For RNA analysis, 2-mL samples from time points were subjected to hot phenol extraction to isolate total RNA. After reverse transcription (Vazyme, R333-01), RNA was quantified by RT-qPCR (Takara Bio, RR820A) using *ACT1* mRNA as an internal control.

### Gene Cloning and Expression in Yeast

The pRS shuttle vector, containing an autonomously replicating sequence and a centromeric sequence, was used to leverage the endogenous replication and chromosome segregation machinery in yeast. This vector also incorporates the G418 resistance gene (KanMX) to allow selection. Vectors were introduced into yeast cells by electroporation.

For *CCW22* structure disruption, the *CCW22* ORF plus 1,000 nucleotides upstream of the start codon, containing its endogenous promoter, was cloned into the pRS vector as a wildtype control. Mutations that either disrupted or restored the *CCW22* structure were introduced into this vector. G418-resistant clones were induced to undergo meiosis. Yeast culture (2 ml) was collected for protein extraction and western blot analysis.

For meiosis-specific overexpression of Ded1, we initially assessed the promoters (1,000 bp sequence upstream start codon) of 10 candidate genes using a GFP reporter system. Yeast cultures (200 μL) from the same batch were sampled at various stages during meiosis, fixed with 3.7% formaldehyde, and the GFP fluorescence intensity was measured using a microplate reader. Among the candidates, the *CDC5* promoter showed the most consistent increase in GFP intensity post-Ndt80 induction (Fig. S4B), and this promoter was used for Ded1 overexpression. The coding region of *DED1* was cloned into the pRS shuttle vector under the control of the *CDC5* promoter. This vector was introduced into yeast cells to assess the effects of Ded1 overexpression.

### DMS-MaP Data Processing

The SK1 genome sequence and annotation were obtained from a previously published dataset^15^. Adaptor sequences were removed using cutadapt^41^, and rRNA reads were depleted using a bowtie2-based alignment^42^ against rDNA sequences. For mRNA abundance quantification, DMSO library reads were mapped to the SK1 genome using STAR aligner^43^, with assigned reads counted to each gene in a strand-specific manner using featureCounts^44^. Finally, read counts were normalized by a TPM method.

For DMS-MaP, antisense reads were removed by bowtie2-based alignment against the assembled transcriptome. The mutation counts and sequencing depth at single-nucleotide resolution were parsed by ShapeMapper2^22^ using filtered fastq files as input. For stagespecific DMS reactivity, data from two replicates were merged to calculate the mutation rate, which was further normalized per transcript by the mean mutation rate of the 90-95% most reactive bases. For overall structures, DMS reactivity was calculated in a similar manner, but by pooling data from all stages.

### Protein Mass Spectrometry Data Processing

Raw mass spectrometry data were quantified using MaxQuant^45^ against the yeast proteome reference from UniProt (strain ATCC 204508 / S288c). The false discovery rate was set at 1% at both the peptide and protein levels. Only proteins identified with at least one ‘unique/razor peptide’ in all samples were used for subsequent normalization. Total TMT intensity in each sample was scaled to 10^6^ (total normalization) as described previously^18^.

### Ribosome Flux and Ribosome Footprint Analyses

To quantify ribosome flux across ORFs, we divided each coding region into 50-nucleotide windows, starting from the 30th nucleotide downstream of the start codon and ending 30 nucleotides upstream of the stop codon. Ribosome footprint (A-site) counts were summed within each window, and the Gini indices were calculated for windows containing more than 20 adenosine or cytosine nucleotides. The window with the highest Gini index was defined as position 0, and ribosome flux was evaluated in the immediately adjacent upstream (–1) and downstream (+1) windows. For uORF analysis, ribosome footprints within the –100 to +900 nucleotide region relative to the annotated start codon were normalized to a total of 1000 counts per transcript.

### RNA Secondary Structure Modeling

DMS reactivities for adenine and cytosine with sequencing depths over 350 in both DMS-treated and DMSO-treated samples were used for structural modeling. The consensus structure model and base-pairing probabilities were predicted using SuperFold^4^, considering long-range interactions (maxPairingDist = 1000). Structural models were visualized with vaRNA^46^, and base-pair probabilities were plotted using R4RNA^47^.

### Meta-gene Analysis

The average DMS reactivity was calculated for adenine and cytosine with sequencing depths over 350, based on their relative positions to the first nucleotide of the start codon or the last nucleotide of the stop codon.

### Hierarchical Clustering

For RNA-seq and ribosome profiling data, data for genes with more than an 8-fold change (*P* < 0.05) in RNA abundance or ribosome footprint across all time points were log-transformed and used for subsequent clustering analysis. For quantitative proteomics data, data for proteins that changed by more than 1.2 fold (*P* < 0.05) across four sampled time points were used for clustering analysis. For co-hierarchical clustering of RNA helicase levels at the translation and protein levels, ribosome footprint and mass spectrometry data were first z-scored and then combined for clustering analysis. Hierarchical clustering was performed using Cluster 3.0^48^ with average clustering based on uncentered Pearson correlation. Clustering data were visualized using Java Treeview^49^.

### Differential Expression Analysis and function Analysis

Fold changes in protein abundance and p-values were calculated using edgeR^50^. Proteins with a fold change of more than 1.2 (*P* < 0.05) were defined as differentially expressed. Gene Ontology enrichment analysis was conducted using g:Profiler^51^. Gene Set Enrichment Analysis was performed using GSEApy^52^, with ranked protein expression from a predefined TUMs gene set as input.

### Statistics and Reproducibility

Unless specified, statistical tests were performed using the Scipy v.1.13 package in Python. The correlation between Gini index and TE was determined based on Spearman’s correlation coefficient. Comparisons between correlation coefficients shown in Fig. 2D were performed using the independent samples *t*-test. Comparisons between Gini indices were performed using the Mann-Whitney U test. Gene overlap analyses between clusters shown in Fig. 3D were performed using the hypergeometric test. Multiple testing corrections were applied using the False Discovery rate method to adjust p-values. The numbers of data points for each analysis are indicated in the figure legends. Unless specified, sequencing and mass spectrometry experiments were performed in two biological replicates, and other experiments were repeated at least three times with consistent results.

## SUPPLEMENTAL FIGURE TITLES AND LEGENDS

**Figure S1.**
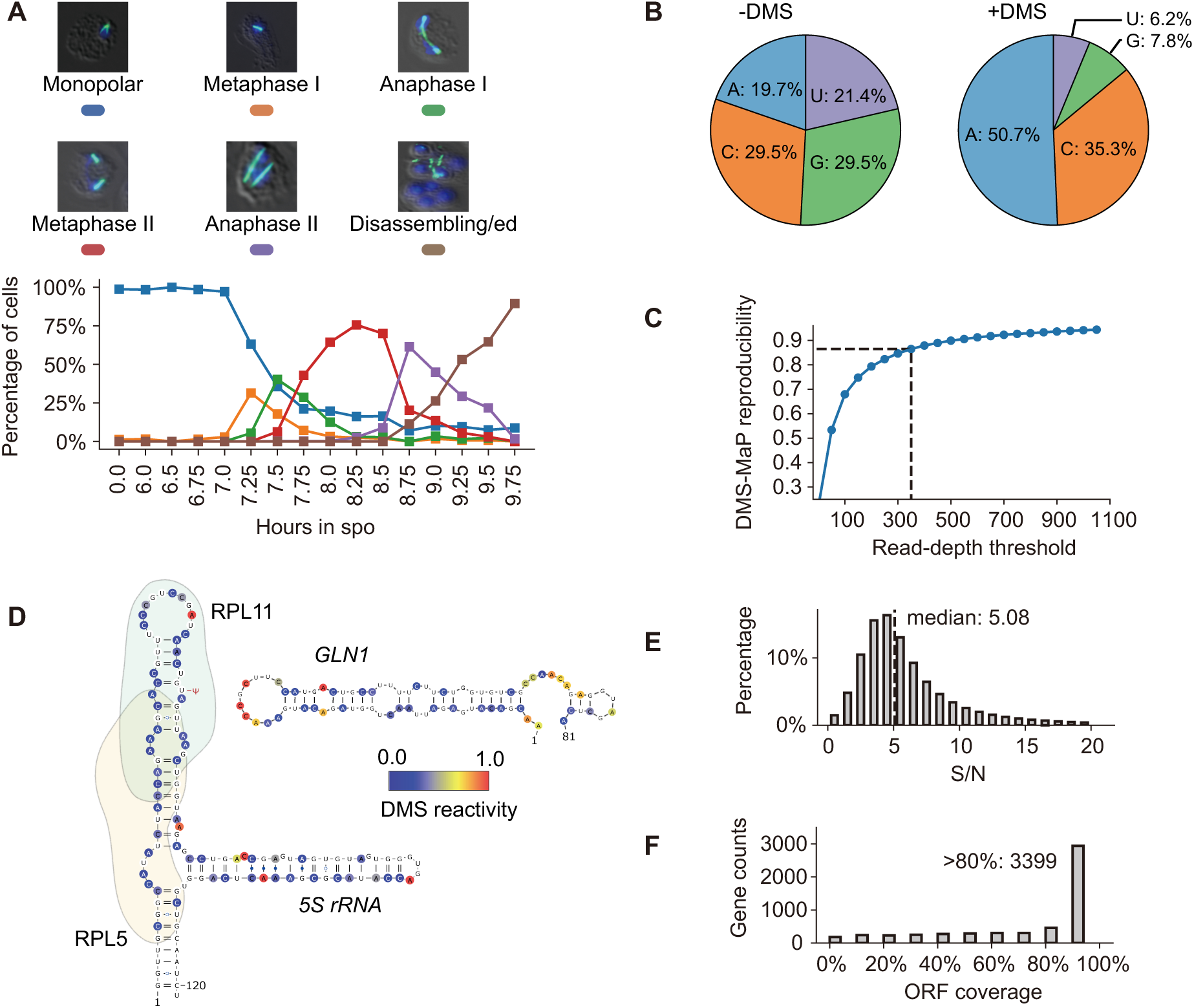
Timepoint staging and DMS-MaP data quality control, related to Figure 1. (A) Upper: Spindle morphologies at different stages (blue, DAPI; green, anti-tubulin). Lower: Percentages of cells at different stages at sampled time points during synchronous meiosis. (B) Mutation rates of four nucleotides as detected by DMS-MaP. +DMS denotes libraries treated with DMS, and -DMS indicates controls treated only with DMSO. (C) Pearson correlation coefficients of DMS reactivities between two replicates at various read-depth thresholds. The dotted line marks the correlation at a 350× read-depth threshold. (D) Data-directed *in vivo* structural model of 5S rRNA, highlighting the binding sites for ribosomal proteins Rpl5 and Rpl11. (E) Distribution of signal-to-noise ratios in DMS reactivity data for nucleotides with read-depths greater than 350. (F) Distribution of ORF coverage for genes with resolved overall structures.

**Figure S2.**
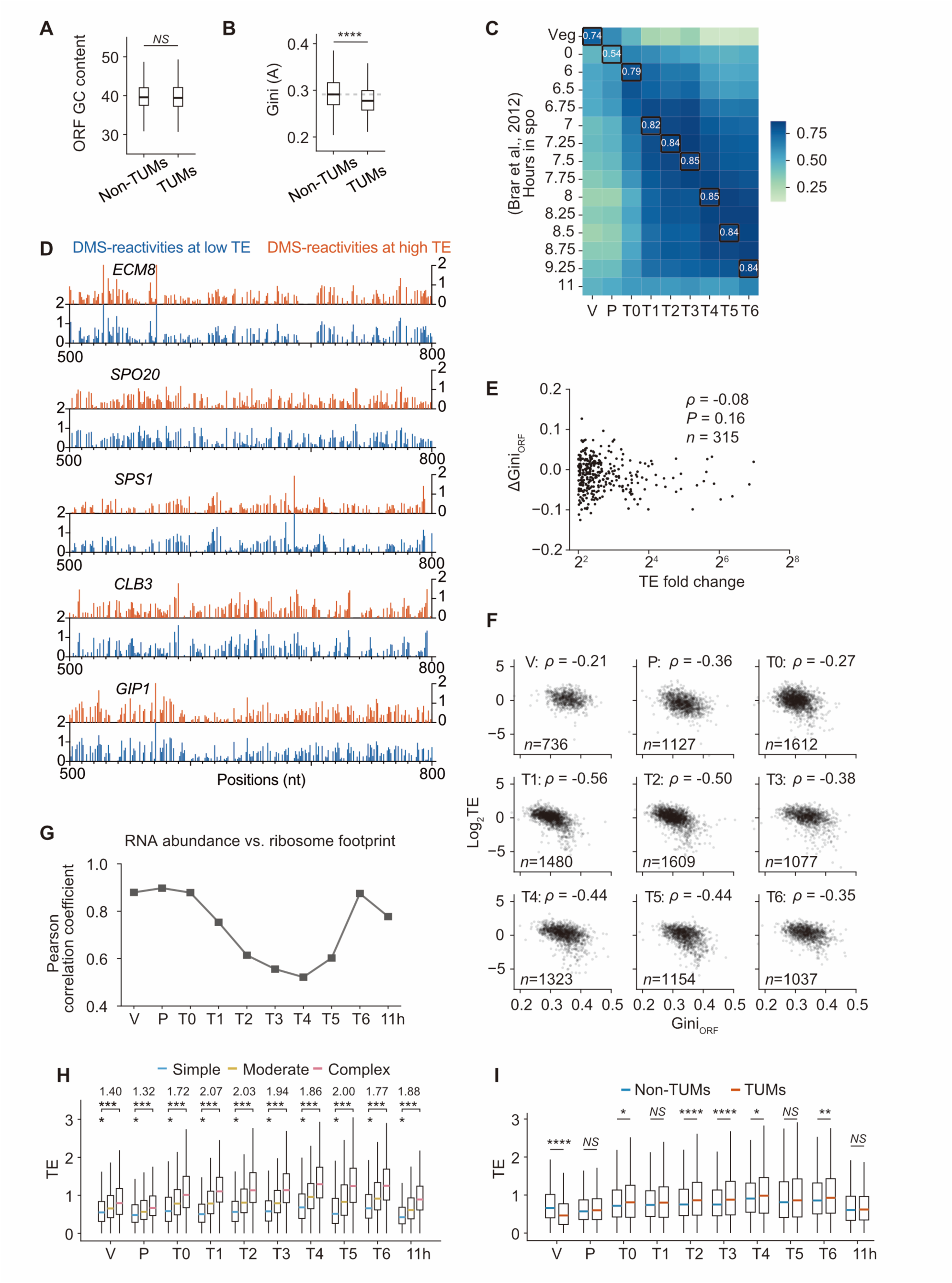
Relationship between mRNA structural complexity and translation efficiency during meiosis, related to Figure 2. (A) Comparison of GC contents between TUMs and non-TUMs. (B) Comparison of Gini indices measured using only data for adenosine between TUMs and non-TUMs. (C) Alignment of sampling time points with the ribosome profiling dataset from a previous study^15^ through RNA-seq profiles. (D) Comparison of DMS reactivities within 300-nucleotide coding regions of five transcripts that have high or low TEs depending on meiotic stages. Fold changes in TE: *ECM8*, 106fold; *SPO20*, 41-fold; *SPS1*, 25-fold; *CLB3*, 20-fold; *GIP1*, 15-fold. (E) Relationship between changes in TEs and Gini indices for ORFs of transcripts with a TE fold change greater than 4. (F) Spearman correlation analysis between TEs and Gini_ORF_ indices at matched time points, related to Fig. 2D. (G) Pearson correlation analysis between mRNA abundance and translation across all time points. (H) Distribution of TEs for mRNA groups with different structural complexities across meiosis. (I) Distribution of TEs for TUMs and non-TUMs across meiosis.

**Figure S3.**
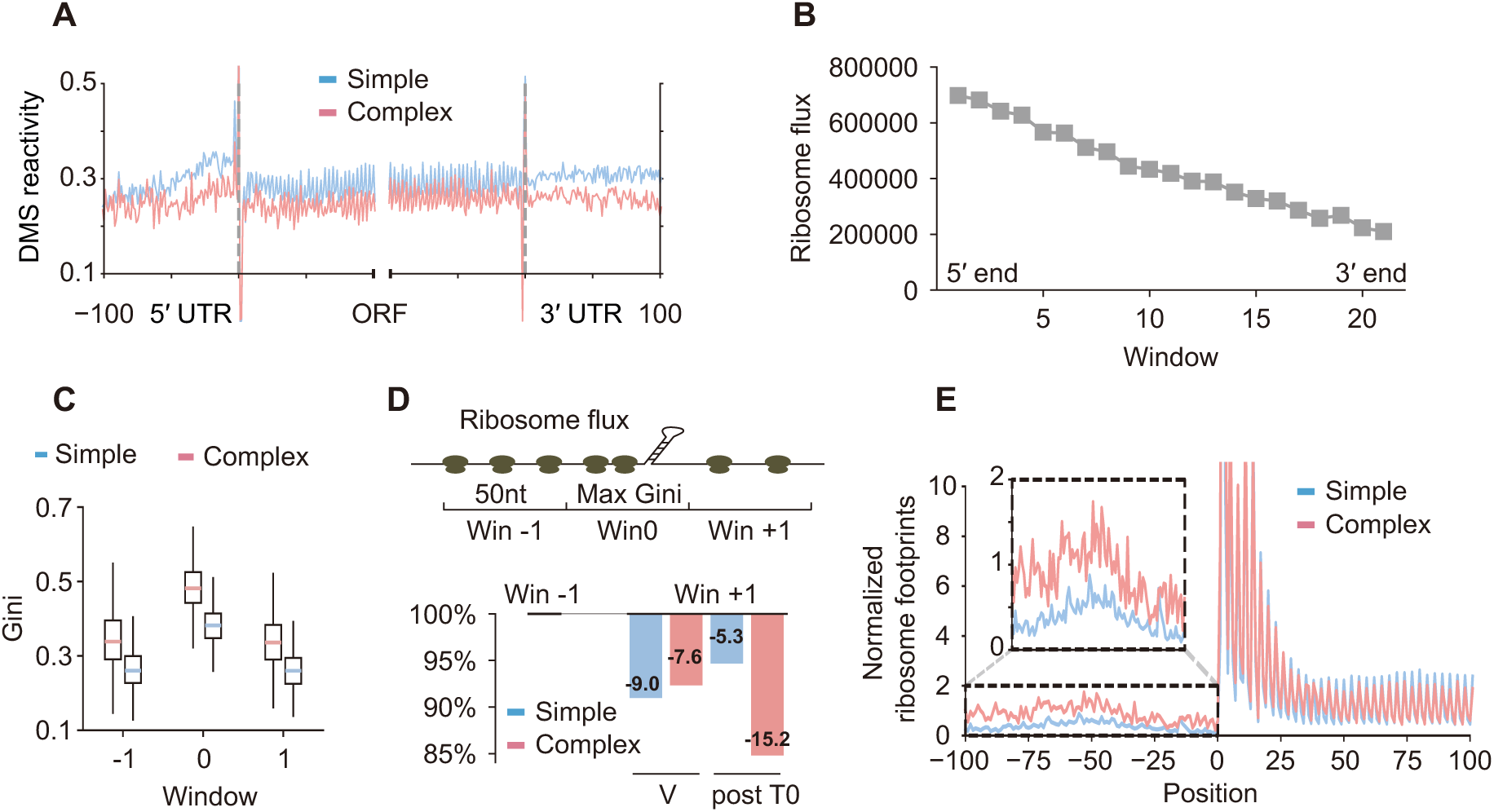
RNA structure impedes ribosome processivity in 5′ UTR and ORF regions during meiosis. (A) Meta-gene analysis of DMS reactivities in Simple and Complex mRNA groups. (B) Ribosome flux from the 1st to the 20th windows (50-nt) from the 5′ end to the 3′ end of mRNA ORF regions, related to Fig. 2G. (C) Gini indices of the windows with the highest Gini index (window 0) and their adjacent windows (window −1 and +1) in Simple and Complex mRNA groups. The window with the maximal Gini index was designated as window 0. (D) Upper: Schematic of regions used for ORF ribosome flux analysis. Ribosome flux was defined as the sum of ribosome footprint reads (A site) within each 50-nt window. Ribosome flux in the upstream window (−1) was used for scaling (to 100%) for comparison. Lower: Percentages of the decrease in ribosome flux from window (−1) to window (+1). (E) Normalized ribosome footprints on uORFs for Simple and Complex TUMs. For each gene, ribosome footprints (A site) from −100 to +900 positions relative to the start codon were normalized to a total of 1,000. For each group, the distribution of ribosome footprints was averaged by position.

**Figure S4.**
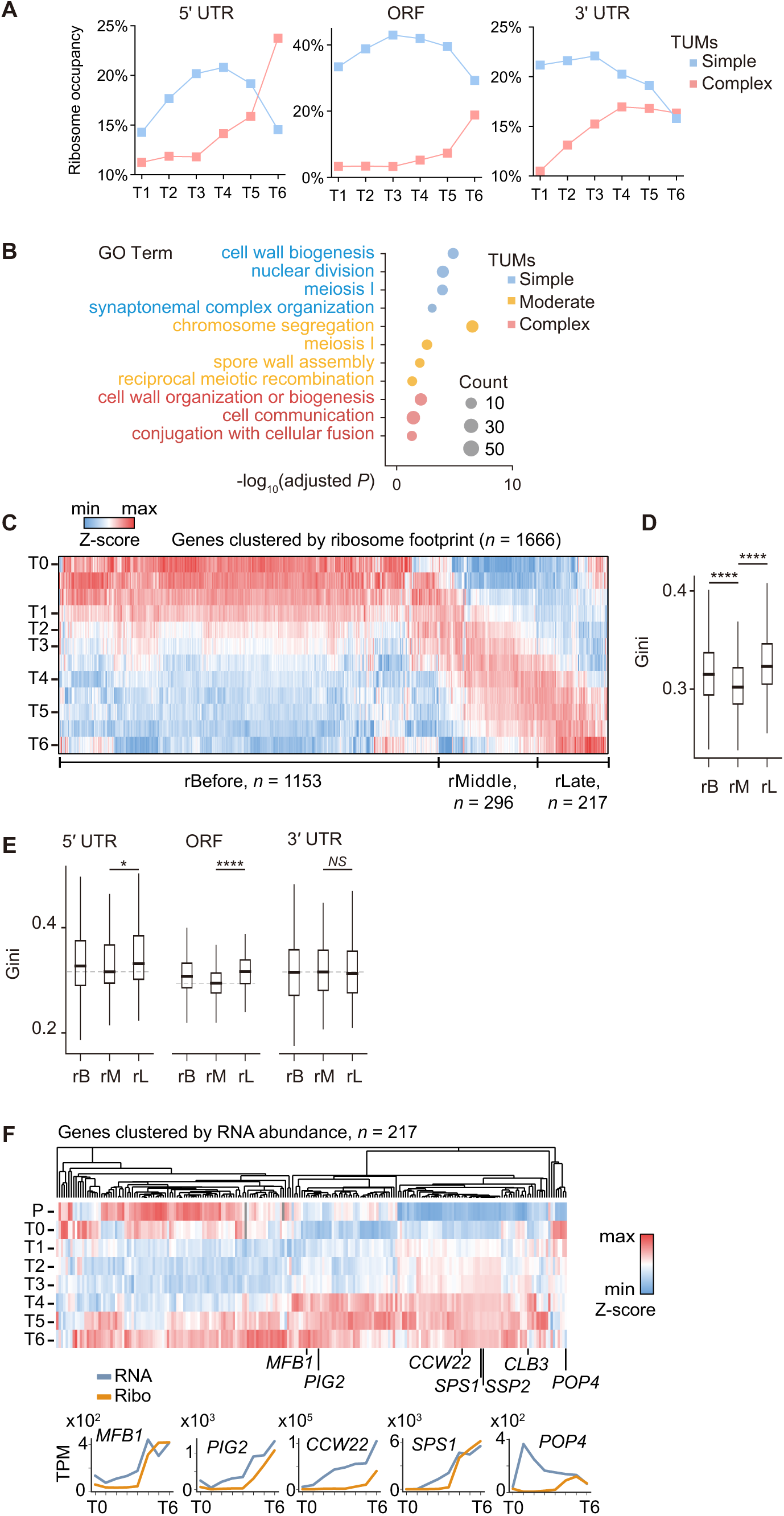
RNA structure-related temporal regulation of translation during meiosis, related to Figure 3. (A) Dynamic patterns of ribosome occupancy among the Simple and Complex mRNA groups classified based on structural complexities in 5′ UTR, ORF, and 3′ UTR regions, related to Fig. 3A. (B) Gene Ontology enrichment analysis of three mRNA groups with different structural complexities, related to Fig. 3A. (C) Hierarchical clustering of ribosome footprint for genes (*n* = 1,666) exhibiting more than an 8-fold change (*P* < 0.05) in TPM across meiosis, related to Fig. 3E. (D) Comparison of Gini indices among three ribosome profiling groups: before meiosis (rBefore, *n* = 930); during mid-meiosis (rMiddle, *n* = 268), and near meiotic exit (rLate, *n* = 188). (E) Comparison of Gini indices among ribosomal profiling clusters (rB, before; rM, mid-meiosis; rL, late) across the 5′ UTR (rB, *n* = 331; rM, *n* = 184; rL, *n* = 121), ORF (rB, *n* = 930; rM, *n* = 268; rL, *n* = 188), and 3′ UTR (rB, *n* = 751; rM, *n* = 227; rL, *n* = 143). Only UTRs with coverage of more than 20 adenines and cytosines were included. (F) Upper: Hierarchical clustering of RNA abundance profiles (TPM) for genes in the rLate group. Lower: RNA abundances and ribosome footprints throughout meiosis for five genes.

**Figure S5.**
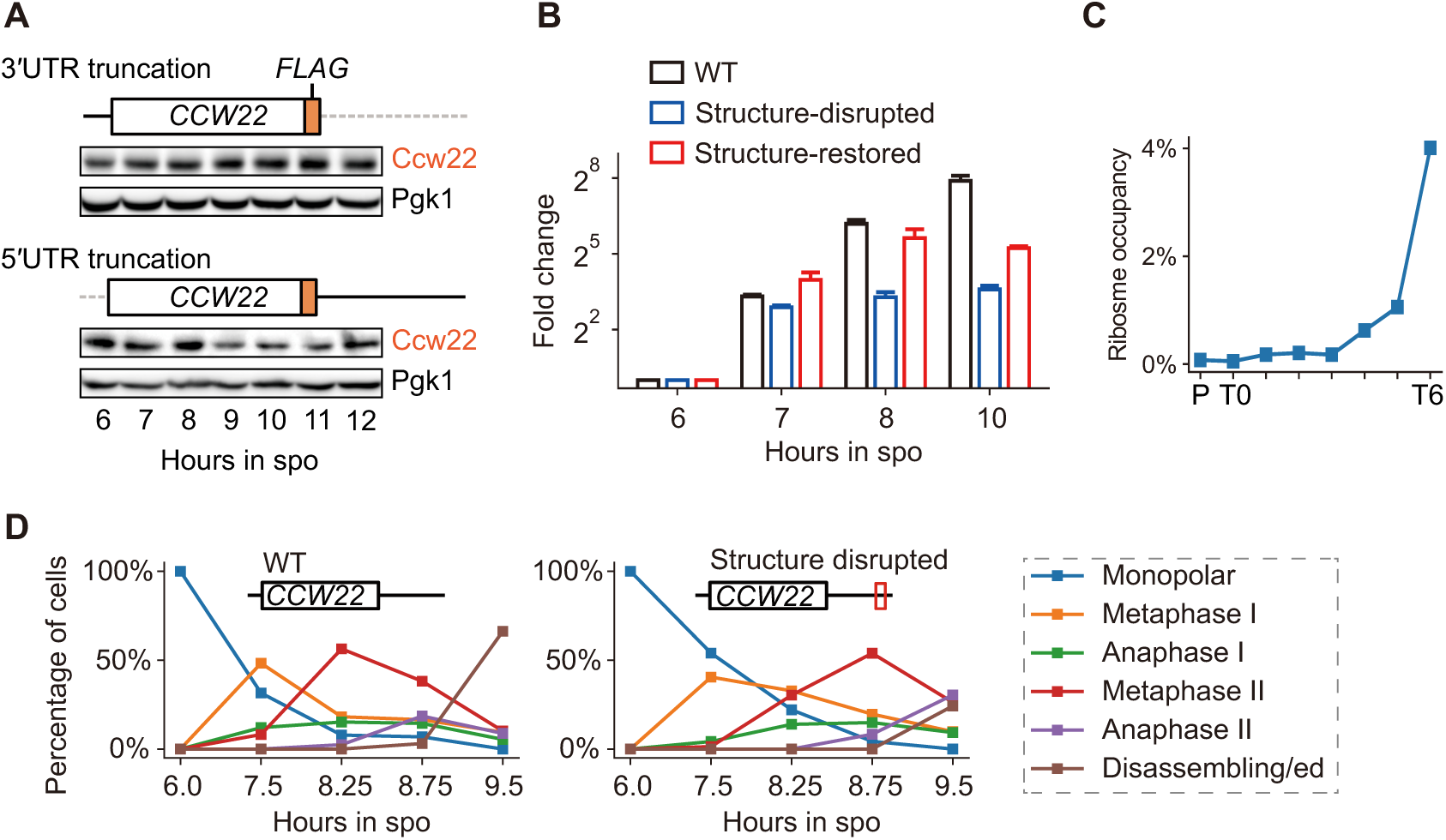
Quantification of CCW22 reporter mRNA and protein levels, and its effects on meiosis, related to Figure 3. (A) Western blot analysis showing Ccw22 expression in yeast strains that express *CCW22* mRNAs with 3′ UTR or 5′ UTR truncation. (B) Abundances of *CCW22* mRNA, quantified by RT-qPCR, expressed from wild-type, structure-disrupted, and structure-restored vectors, related to Fig. 3I. Note that compensatory mutations altered the Kozak sequence, which may have slightly reduced translation efficiency from this construct; this effect does not influence our conclusions about expression timing. (C) Ribosome occupancy of *CCW22* mRNA across meiosis. (D) Percentage of cells at each meiotic stage in yeast strains harboring vectors for expression of wild-type *CCW22* or *CCW22* mRNA with mutation of the 3′ UTR.

**Figure S6.**
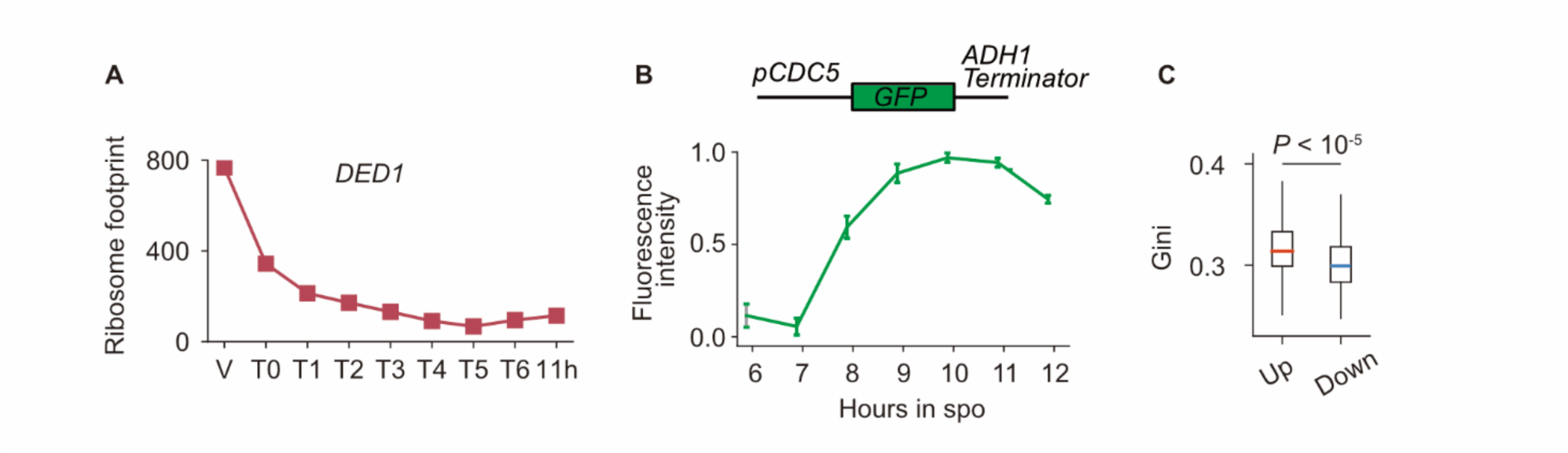
Analysis of *DED1* expression and effects of Ded1 overexpression, related to Figure 4. (A) Ribosome footprint for *DED1* mRNA across meiosis. (B) GFP intensity driven by the *CDC5* promoter. (C) Comparison of Gini indices for upregulated and downregulated genes in cells that overexpress Ded1.

## Supplemental information

Table S1. DMS-MaP sequencing depth for each stage.

Table S2. Quantitative data including RNA abundance, RNA structural complexity, matched ribosome footprint, and protein abundance across meiosis.

Table S3. Statistics of stage-specific structures.

Table S4. Gene Ontology Enrichment Analysis results, related to Fig. S4B.

Table S5. Publication enrichment from the Saccharomyces Genome Database.

Table S6. The classification by structural complexity of transcripts with early and delayed translation timing, related to Fig. S4C and S4D.

Table S7. Structure complexity of 5′UTR, ORF, and 3′UTR regions, related to Fig. 4D, S4A and S4E.

Table S8. RNA helicase expression at translation and protein levels, related to Fig. 4A, 4B and S6A.

Table S9. Differential gene expression analysis upon Ded1 overexpression, related to Figure 4D and 4F.

## References

1. Dolgin, E. (2015). The Elaborate Architecture of RNA. Nature 523, 398–399. 10.1038/523398a.

2. Watts, J.M., Dang, K.K., Gorelick, R.J., Leonard, C.W., Bess Jr, J.W., Swanstrom, R., Burch, C.L., and Weeks, K.M. (2009). Architecture and secondary structure of an entire HIV-1 RNA genome. Nature 460, 711–716. 10.1038/nature08237.

3. Burkhardt, D.H., Rouskin, S., Zhang, Y., Li, G.-W., Weissman, J.S., and Gross, C.A. (2017). Operon mRNAs are organized into ORF-centric structures that predict translation efficiency. eLife 6, e22037. 10.7554/eLife.22037.

4. Mustoe, A.M., Busan, S., Rice, G.M., Hajdin, C.E., Peterson, B.K., Ruda, V.M., Kubica, N., Nutiu, R., Baryza, J.L., and Weeks, K.M. (2018). Pervasive Regulatory Functions of mRNA Structure Revealed by High-Resolution SHAPE Probing. Cell 173, 181–195.e18. 10.1016/j.cell.2018.02.034.

5. Rouskin, S., Zubradt, M., Washietl, S., Kellis, M., and Weissman, J.S. (2014). Genomewide probing of RNA structure reveals active unfolding of mRNA structures in vivo. Nature 505, 701–705. 10.1038/nature12894.

6. Ding, Y., Tang, Y., Kwok, C.K., Zhang, Y., Bevilacqua, P.C., and Assmann, S.M. (2014). In vivo genome-wide profiling of RNA secondary structure reveals novel regulatory features. Nature 505, 696–700. 10.1038/nature12756.

7. Wan, Y., Qu, K., Zhang, Q.C., Flynn, R.A., Manor, O., Ouyang, Z., Zhang, J., Spitale, R.C., Snyder, M.P., Segal, E., et al. (2014). Landscape and variation of RNA secondary structure across the human transcriptome. Nature 505, 706–709. 10.1038/nature12946.

8. Sun, L., Fazal, F.M., Li, P., Broughton, J.P., Lee, B., Tang, L., Huang, W., Kool, E.T., Chang, H.Y., and Zhang, Q.C. (2019). RNA structure maps across mammalian cellular compartments. Nat Struct Mol Biol 26, 322–330. 10.1038/s41594-019-0200-7.

9. Beaudoin, J.-D., Novoa, E.M., Vejnar, C.E., Yartseva, V., Takacs, C.M., Kellis, M., and Giraldez, A.J. (2018). Analyses of mRNA structure dynamics identify embryonic gene regulatory programs. Nat Struct Mol Biol 25, 677–686. 10.1038/s41594-018-0091-z.

10. Xiang, Y., Huang, W., Tan, L., Chen, T., He, Y., Irving, P.S., Weeks, K.M., Zhang, Q.C., and Dong, X. (2023). Pervasive downstream RNA hairpins dynamically dictate startcodon selection. Nature 621, 423–430. 10.1038/s41586-023-06500-y.

11. Metelev, M., Lundin, E., Volkov, I.L., Gynnå, A.H., Elf, J., and Johansson, M. (2022). Direct measurements of mRNA translation kinetics in living cells. Nat Commun 13, 1852. 10.1038/s41467-022-29515-x.

12. Halstead, J.M., Lionnet, T., Wilbertz, J.H., Wippich, F., Ephrussi, A., Singer, R.H., and Chao, J.A. (2015). An RNA biosensor for imaging the first round of translation from single cells to living animals. Science 347, 1367–1671. 10.1126/science.aaa3380.

13. Wu, B., Eliscovich, C., Yoon, Y.J., and Singer, R.H. (2016). Translation dynamics of single mRNAs in live cells and neurons. Science 352, 1430–1435. 10.1126/science.aaf1084.

14. Chu, S., DeRisi, J., Eisen, M., Mulholland, J., Botstein, D., Brown, P.O., and Herskowitz, I. (1998). The Transcriptional Program of Sporulation in Budding Yeast. Science 282, 699–705. 10.1126/science.282.5389.699.

15. Brar, G.A., Yassour, M., Friedman, N., Regev, A., Ingolia, N.T., and Weissman, J.S. (2012). High-Resolution View of the Yeast Meiotic Program Revealed by Ribosome Profiling. Science 335, 552–557. 10.1126/science.1215110.

16. Berchowitz, L.E., Kabachinski, G., Walker, M.R., Carlile, T.M., Gilbert, W.V., Schwartz, T.U., and Amon, A. (2015). Regulated Formation of an Amyloid-like Translational Repressor Governs Gametogenesis. Cell 163, 406–418. 10.1016/j.cell.2015.08.060.

17. Guenther, U.-P., Weinberg, D.E., Zubradt, M.M., Tedeschi, F.A., Stawicki, B.N., Zagore, L.L., Brar, G.A., Licatalosi, D.D., Bartel, D.P., Weissman, J.S., et al. (2018). The helicase Ded1p controls use of near-cognate translation initiation codons in 5′UTRs. Nature 559, 130–134. 10.1038/s41586-018-0258-0.

18. Cheng, Z., Otto, G.M., Powers, E.N., Keskin, A., Mertins, P., Carr, S.A., Jovanovic, M., and Brar, G.A. (2018). Pervasive, Coordinated Protein-Level Changes Driven by Transcript Isoform Switching during Meiosis. Cell 172, 910–923.e16. 10.1016/j.cell.2018.01.035.

19. Carlile, T.M., and Amon, A. (2008). Meiosis I Is Established through Division-Specific Translational Control of a Cyclin. Cell 133, 280–291. 10.1016/j.cell.2008.02.032.

20. Keeling, J.W.P., and Miller, R.K. (2011). Indirect Immunofluorescence for Monitoring Spindle Assembly and Disassembly in Yeast. In Cell Cycle Checkpoints: Methods and Protocols Methods in Molecular Biology., W. X. name>Li, ed. (Humana Press), pp. 231–244. 10.1007/978-1-61779-273-1_17.

21. Zubradt, M., Gupta, P., Persad, S., Lambowitz, A.M., Weissman, J.S., and Rouskin, S. (2017). DMS-MaPseq for genome-wide or targeted RNA structure probing in vivo. Nat Methods 14, 75–82. 10.1038/nmeth.4057.

22. Busan, S., and Weeks, K.M. (2018). Accurate detection of chemical modifications in RNA by mutational profiling (MaP) with ShapeMapper 2. RNA 24, 143–148. 10.1261/rna.061945.117.

23. Madru, C., Lebaron, S., Blaud, M., Delbos, L., Pipoli, J., Pasmant, E., Réty, S., and Leulliot, N. (2015). Chaperoning 5S RNA assembly. Genes Dev. 29, 1432–1446. 10.1101/gad.260349.115.

24. Rüegsegger, U., Leber, J.H., and Walter, P. (2001). Block of HAC1 mRNA Translation by Long-Range Base Pairing Is Released by Cytoplasmic Splicing upon Induction of the Unfolded Protein Response. Cell 107, 103–114. 10.1016/S0092-8674(01)00505-0.

25. Aragón, T., van Anken, E., Pincus, D., Serafimova, I.M., Korennykh, A.V., Rubio, C.A., and Walter, P. (2009). Messenger RNA targeting to endoplasmic reticulum stress signalling sites. Nature 457, 736–740. 10.1038/nature07641.

26. Rouskin, S., Zubradt, M., Washietl, S., Kellis, M., and Weissman, J.S. (2014). Genomewide probing of RNA structure reveals active unfolding of mRNA structures in vivo. Nature 505, 701–705. 10.1038/nature12894.

27. Chu, S., and Herskowitz, I. (1998). Gametogenesis in Yeast Is Regulated by a Transcriptional Cascade Dependent on Ndt80. Molecular Cell 1, 685–696. 10.1016/S1097-2765(00)80068-4.

28. Bao, C., Loerch, S., Ling, C., Korostelev, A.A., Grigorieff, N., and Ermolenko, D.N. (2020). mRNA stem-loops can pause the ribosome by hindering A-site tRNA binding. eLife 9, e55799. 10.7554/eLife.55799.

29. Jin, L., Zhang, K., Xu, Y., Sternglanz, R., and Neiman, A.M. (2015). Sequestration of mRNAs Modulates the Timing of Translation during Meiosis in Budding Yeast. Mol Cell Biol 35, 3448–3458. 10.1128/MCB.00189-15.

30. Bleichert, F., and Baserga, S.J. (2007). The Long Unwinding Road of RNA Helicases. Molecular Cell 27, 339–352. 10.1016/j.molcel.2007.07.014.

31. Li, L., Krasnykov, K., Homolka, D., Gos, P., Mendel, M., Fish, R.J., Pandey, R.R., and Pillai, R.S. (2022). The XRN1-regulated RNA helicase activity of YTHDC2 ensures mouse fertility independently of m6A recognition. Molecular Cell 82, 1678–1690.e12. 10.1016/j.molcel.2022.02.034.

32. Zhang, K., Zhang, T., Zhang, Y., Yuan, J., Tang, X., Zhang, C., Yin, Q., Zhang, Y., and Tong, M.-H. (2023). DNA/RNA helicase DHX36 is required for late stages of spermatogenesis. Journal of Molecular Cell Biology 14, mjac069. 10.1093/jmcb/mjac069.

33. Tsai-Morris, C.-H., Sheng, Y., Lee, E., Lei, K.-J., and Dufau, M.L. (2004). Gonadotropin-regulated testicular RNA helicase (GRTH/Ddx25) is essential for spermatid development and completion of spermatogenesis. Proceedings of the National Academy of Sciences 101, 6373–6378. 10.1073/pnas.0401855101.

34. Gupta, N., Lorsch, J.R., and Hinnebusch, A.G. (2018). Yeast Ded1 promotes 48S translation pre-initiation complex assembly in an mRNA-specific and eIF4F-dependent manner. Elife 7, e38892. 10.7554/eLife.38892.

35. Powers, E.N., Kuwayama, N., Sousa, C., Reynaud, K., Jovanovic, M., Ingolia, N.T., and Brar, G.A. (2024). Dbp1 is a low performance paralog of RNA helicase Ded1 that drives impaired translation and heat stress response. bioRxiv, 2024.01.12.575095. 10.1101/2024.01.12.575095.

36. Liu, X., Yao, Z., Jin, M., Namkoong, S., Yin, Z., Lee, J.H., and Klionsky, D.J. (2019). Dhh1 promotes autophagy-related protein translation during nitrogen starvation. PLoS Biol 17, e3000219. 10.1371/journal.pbio.3000219.

37. Sen, N.D., Zhou, F., Ingolia, N.T., and Hinnebusch, A.G. (2015). Genome-wide analysis of translational efficiency reveals distinct but overlapping functions of yeast DEAD-box RNA helicases Ded1 and eIF4A. Genome Res 25, 1196–1205. 10.1101/gr.191601.115.

38. Bohnsack, K.E., Yi, S., Venus, S., Jankowsky, E., and Bohnsack, M.T. (2023). Cellular functions of eukaryotic RNA helicases and their links to human diseases. Nat Rev Mol Cell Biol 24, 749–769. 10.1038/s41580-023-00628-5.

39. Ares, M. (2012). Isolation of total RNA from yeast cell cultures. Cold Spring Harb Protoc 2012, 1082–1086. 10.1101/pdb.prot071456.

40. Smola, M.J., Rice, G.M., Busan, S., Siegfried, N.A., and Weeks, K.M. (2015). Selective 2′-hydroxyl acylation analyzed by primer extension and mutational profiling (SHAPEMaP) for direct, versatile and accurate RNA structure analysis. Nat Protoc 10, 1643– 1669. 10.1038/nprot.2015.103.

41. Martin, M. (2011). Cutadapt removes adapter sequences from high-throughput sequencing reads. EMBnet.journal 17, 10–12. 10.14806/ej.17.1.200.

42. Langmead, B., and Salzberg, S.L. (2012). Fast gapped-read alignment with Bowtie 2. Nat Methods 9, 357–359. 10.1038/nmeth.1923.

43. Dobin, A., Davis, C.A., Schlesinger, F., Drenkow, J., Zaleski, C., Jha, S., Batut, P., Chaisson, M., and Gingeras, T.R. (2013). STAR: ultrafast universal RNA-seq aligner. Bioinformatics 29, 15–21. 10.1093/bioinformatics/bts635.

44. Liao, Y., Smyth, G.K., and Shi, W. (2014). featureCounts: an efficient general purpose program for assigning sequence reads to genomic features. Bioinformatics 30, 923– 930. 10.1093/bioinformatics/btt656.

45. Tyanova, S., Temu, T., and Cox, J. (2016). The MaxQuant computational platform for mass spectrometry-based shotgun proteomics. Nat Protoc 11, 2301–2319. 10.1038/nprot.2016.136.

46. Darty, K., Denise, A., and Ponty, Y. (2009). VARNA: Interactive drawing and editing of the RNA secondary structure. Bioinformatics 25, 1974–1975. 10.1093/bioinformatics/btp250.

47. Lai, D., Proctor, J.R., Zhu, J.Y.A., and Meyer, I.M. (2012). R-CHIE: a web server and R package for visualizing RNA secondary structures. Nucleic Acids Res 40, e95. 10.1093/nar/gks241.

48. de Hoon, M.J.L., Imoto, S., Nolan, J., and Miyano, S. (2004). Open source clustering software. Bioinformatics 20, 1453–1454. 10.1093/bioinformatics/bth078.

49. Saldanha, A.J. (2004). Java Treeview--extensible visualization of microarray data. Bioinformatics 20, 3246–3248. 10.1093/bioinformatics/bth349.

50. Robinson, M.D., McCarthy, D.J., and Smyth, G.K. (2010). edgeR: a Bioconductor package for differential expression analysis of digital gene expression data. Bioinformatics 26, 139–140. 10.1093/bioinformatics/btp616.

51. Reimand, J., Kull, M., Peterson, H., Hansen, J., and Vilo, J. (2007). g:Profiler—a webbased toolset for functional profiling of gene lists from large-scale experiments. Nucleic Acids Res 35, W193–W200. 10.1093/nar/gkm226.

52. Fang, Z., Liu, X., and Peltz, G. (2023). GSEApy: a comprehensive package for performing gene set enrichment analysis in Python. Bioinformatics 39, btac757. 10.1093/bioinformatics/btac757.

